# The complex history of genome duplication and hybridization in North American gray treefrogs

**DOI:** 10.1101/2020.11.25.398461

**Authors:** William W. Booker, H. Carl Gerhardt, Alan R. Lemmon, Margaret B. Ptacek, Alyssa T. B. Hassinger, Johannes Schul, Emily Moriarty Lemmon

## Abstract

Polyploid speciation has played an important role in evolutionary history across the tree of life, yet there remain large gaps in our understanding of how polyploid species form and persist. While systematic studies have been conducted in numerous polyploid complexes, recent advances in sequencing technology have demonstrated that conclusions from data-limited studies may be spurious and misleading. The North American gray treefrog complex, consisting of the diploid *Hyla chrysoscelis* and the tetraploid *Hyla versicolor*, has long been used as a model system in a variety of biological fields, yet all taxonomic studies to date were conducted with only a few loci from nuclear and mitochondrial genomes. Here, we utilized anchored hybrid enrichment and high-throughput sequencing to capture hundreds of loci along with whole mitochondrial genomes to investigate the evolutionary history of this complex. We used several phylogenetic and population genetic methods, including coalescent simulations and testing of polyploid speciation models with Approximate Bayesian Computation (ABC), to determine that H. versicolor was most likely formed via autopolyploidization from a now extinct lineage of H. chrysoscelis. We also uncovered evidence of significant hybridization between diploids and tetraploids where they co-occur, and show that historical hybridization between these groups led to the re-formation of distinct polyploid lineages following the initial whole genome duplication event. Our study indicates that a wide variety of methods and explicit model testing of polyploid histories can greatly facilitate efforts to uncover the evolutionary history of polyploid complexes.

## Introduction

Hybridization is universal. As the genomic and taxonomic breadth of systematic studies continues to increase, the tree of life less resembles a series of simple bifurcations and instead becomes better defined by a complex network of interactions, exchanges, and rearrangements. Understanding how to disentangle these networks, and how they translate into the formation of distinct and identifiable species, however, remains elusive.

Polyploidization represents a unique form of speciation defined by an increase in the number of chromosome sets a species has in comparison to its ancestral taxa. Though unique in mechanism, polyploidy has undoubtedly played a major role in shaping the tree of life. While only common in some animal clades (Mable et al. 2011), polyploidy is well known as a major driver of diversification in plants (Blischak et al. 2018; One Thousand Plant Transcriptomes Initiative 2019), has occurred at least twice in early vertebrate history (Gregory and Mable 2005; Mable et al. 2011), is likely common in prokaryotes and fungi (Albertin and Marullo 2012), and has recently been discovered to play a major role in the diversification of insects (Li et al. 2018).

Whereas allopolyploidy is defined by hybridization, both allopolyploids and autopolyploids frequently hybridize with relatives of different ploidies during and after their formation (Bogart and Bi 2013; Soltis et al. 2014). The genomic and phenotypic consequences of hybridization across ploidies are however largely unknown. Although ongoing research on polyploid evolution continues to produce novel insights, many aspects of polyploidy may be poorly understood because most polyploid research has been conducted in a few model systems (Dufresne et al. 2014; Soltis et al. 2016) In order to further our knowledge of how polyploidization contributes to speciation, a greater number of well-defined systems must be developed.

The first step of defining any study system is estimating the phylogenetic relationships of its taxa. Our ability to infer these relationships in polyploids, however, is impeded by several unique challenges. Because a single genealogy can be explained by several different evolutionary histories, a particularly difficult challenge is determining the specific process that generated a given phylogenetic pattern. For example, recovering evidence of novel alleles in a polyploid could be explained by either allopolyploidy, whereby novel alleles descended from an extinct heterospecific taxon, or by autopolyploidy, where novel alleles originated from an extinct conspecific population. Additionally, the processes of gene conversion and mixed chromosomal inheritance can make it difficult to distinguish potentially unique genomes within a polyploid individual and accurately infer subgenomic phylogenetic relationships (Dufresne et al. 2014). Fortunately, recent advances in methods for collecting genomic data (Lemmon and Lemmon 2013) and analyzing these data (Blischak et al. 2018) provide powerful tools for elucidating complex genomic histories.

As our technological ability to elucidate polyploid evolutionary histories has expanded, so too has our methodology. Perhaps the most powerful among these methods retrace polyploid origins by linking informative characters with whole genome assays (Tennessen et al. 2014; Douglas et al. 2015; Session et al. 2016). For example, Session et al. (2016) sequenced the whole genome of *Xenopus laevis* and used fluorescence *in situ* hybridization (FISH) to link relict transposable elements to individual chromosomes in order to distinguish between allo- vs. autopolyploidy. This method is powerful, however, in many cases such studies are not economically or logistically feasible—especially for non-model species or those with large genome sizes. Conversely, for cases where a study is limited to sequencing only a few loci, Roux and Pannel (2015) outline a more accessible method that estimates the mode of polyploidization as a model parameter through Approximate Bayesian Computation (ABC) by comparing observed population genetic summary statistics to those from simulated sequence data. This method is inherently less informative than a whole genome assay, but at the very least it purports to reliably indicate the modes of polyploidization and chromosomal inheritance as well an estimate the timing of lineage splits and whole genome duplications. Finally, a considerable body of literature has focused on the use of traditional phylogenetic inference for polyploid analysis, elucidating origins from a few to thousands of sequenced loci, which can provide an informative middle ground between the previous two approaches (Holloway et al. 2006; Rousseau-Gueutin et al. 2009; St. Onge et al. 2012; Brassac and Blattner 2015; Gregg et al. 2017). Though these methods alone can be useful, researchers must be careful in the way data are prepared and analyzed since most software and models for conducting phylogenetic inference have been developed for haploid or diploid taxa. Furthermore, any conclusions drawn from a single phylogenetic analysis must be couched with the understanding that many polyploid histories can produce the same phylogenetic signal.

One system in which the mode of polyploidization remains in question is the North American gray treefrog complex, comprised of the diploid *Hyla chrysoscelis*, and tetraploid *H. versicolor* (Le Conte 1825; Johnson 1963; Wasserman 1970; Bogart and Wasserman 1972). These two species are morphologically indistinguishable and are only reliably identified by their karyotype or male acoustic signal (Gerhardt 2005; Holloway et al. 2006). It is important that we understand the evolutionary history of this group, since the gray treefrog system is utilized for addressing key questions several biological subdisciplines, including behavioral ecology, evolution, genetics, and neurobiology (Bogart and Wasserman 1972; Gerhardt 1974, 1978; Ralin and Selander 1979; Storey and Storey 1985; Wells and Taigen 1986; Gerhardt and Doherty 1988; Ptacek et al. 1994; Relyea and Mills 2001; Gerhardt and Huber 2002; Schul and Bush 2002; Endepols et al. 2004; Holloway et al. 2006; Blair 1962; Littlejohn et al. 1960). Of particular relevance to the present study, research on reproductive isolation in this system demonstrates that in addition to causing immediate postzygotic isolation through duplication of chromosomes, polyploidization may promote prezygotic isolation between ploidy levels by observed shifts in acoustic reproductive behaviors caused by the change in cell size, cell number, and chromosome number (Ueda 1993; Keller and Gerhardt 2001; Tucker and Gerhardt 2012). Estimated differences in pulse rate following polyploidization in *H. versicolor* from synthetically created *H. chrysoscelis* triploids and *H. japonica* tetraploids suggest that *H. versicolor* may have been reproductively isolated from *H. chrysoscelis* immediately upon its formation (Keller and Gerhardt 2001). As such, the gray treefrog system may represent a particularly compelling example of speciation immediately following the formation of a polyploid lineage.

Because of the morphological and chromosomal similarity between diploid and tetraploid gray treefrogs, it was long thought that the tetraploid *H. versicolor* arose from an autopolyploid genome duplication event (Wasserman 1970; Ralin and Selander 1979; Danzmann and Bogart 1983; Bogart and Bi 2013; Bogart et al. 2020). The most recent and comprehensive studies to date on the origins of the tetraploid species suggests, however, that there were multiple allopolyploid origins from hybridization of *H. chrysoscelis* with two other extinct species (Ptacek et al. 1994; Holloway et al. 2006). This hypothesis is supported by the presence of nuclear and mitochondrial alleles in tetraploids that do not exist in any *H. chrysoscelis* or its extant diploid relative *H. avivoca i* as well as previously unexplainable nuclear allozymes (Ralin and Selander 1979; Ptacek et al. 1994). Though groundbreaking for its time, the data limitations of Holloway et al. (2006, 3 nuclear loci and 1 mitochondrial locus included) as well as the complex nature of polyploid genomes precluded rigorous testing of alternative hypotheses of polyploid formation. Indeed, as Bogart and Bi (Bogart and Bi 2013) suggest, the data of the Holloway et al. (2006) study are not explicitly incompatible with alternative conclusions such as an autopolyploid origin of *H. versicolor*, and Bogart et al. (2020) showed that hybridization across ploidies of the two species is likely—a condition Holloway et al. (2006) did not consider.

To fully elucidate the evolutionary history of the North American gray treefrog complex and further develop this system for future studies of polyploidization, we built upon the previous work of Ptacek et al. (1994) and Holloway et al. (2006) by utilizing Anchored Hybrid Enrichment (Lemmon et al. 2012) to collect data from hundreds of nuclear loci and complete mitochondrial genomes. Accounting for the complications of polyploid studies, we used phylogenomic, population genetic, and model testing methods to reconstruct the origins of the tetraploid *H. versicolor* with high confidence. In short, the goals of our study were to: (1) ascertain the mode of polyploidization in *H. versicolor*, (2) determine the identity and number of ancestor(s) that gave rise to tetraploid *H. versicolor*, (3) determine the number of independent origins of polyploidy and the timing of any whole genome duplication events, and (4) characterize the population structure and evolutionary history of the complex.

## Materials and Methods

### Sampling

For molecular data, a total of 35 *Hyla versicolor* and 71 *H. chrysoscelis* were sampled, including nearly all samples from the Ptacek et al. (1994) and Holloway et al. (2006) studies (Supp. Table 1). We also included 7 *H. avivoca,* the closest known relative of the gray treefrogs which often hybridizes with the diploid *H. chrysoscelis*, to examine their genomic contribution to this complex (Gerhardt 1974). Finally, 1 *H. andersonii*, 2 *H. arenicolor*, and 1 *H. femoralis* —three closely related species previously assigned to the *H. versicolor* group, but whose exact relationships remain unclear (Blair 1962; Faivovich et al. 2005; Duellman et al. 2016)—were used as outgroup taxa. Field collections were conducted in accordance with appropriate state collection permits and Animal Care and Use Committee Protocols. Treefrogs were collected by hand and either euthanized, dissected, and vouchered for deposition at the Florida Museum of Natural History or toe-clipped and released at their original locality. Tissues (heart, liver, leg muscle, and/or toe clips) were preserved in tissue buffer (20% DMSO, 0.25 M EDTA, salt-saturated) and stored at −80*^◦^*F at Florida State University. Additional samples were obtained via museum loans from the University of Texas at Austin Biodiversity Center (formerly Texas Natural History Collections), and the Illinois Natural History Survey.

**Table 1.**
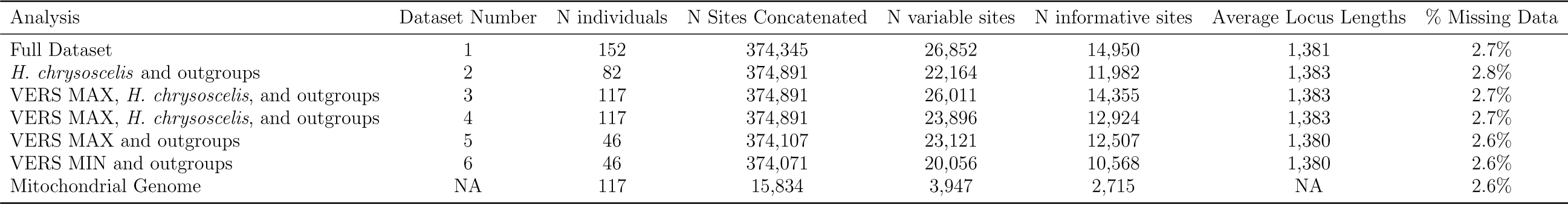
Summary table of genetic data for the analyses from Fig. 4. Note that N sites concatenated is the maximum length of the alignment for that specific analysis.

### Anchored Hybrid Enrichment and Sequencing

Genomic DNA was extracted using an E.Z.N.A. Tissue DNA Kit (Omega Bio-Tek). After extraction, genomic DNA was sonicated to a fragment size of ~300-800 bp using a Covaris E220 Focused-ultrasonicator with Covaris microTUBES. Subsequently, library preparation and indexing were performed on a Beckman-Coulter Biomek FXp liquid-handling robot following a protocol modified from Meyer and Kircher (2010). A size-selection step was also employed after blunt-end repair using SPRI select beads (Beckman-Coulter Inc.; 0.9x ratio of bead to sample volume). Indexed samples were then pooled at equal quantities (16-18 samples per pool), and then each pool was enriched using amphibian-specific probes (Agilent Technologies Custom SureSelect XT kit) described elsewhere (Barrow et al. 2018; Heinicke et al. 2018). Capture probes were designed based on 15x raw genomic data (Illumina 2500 paired-end 150bp) derived from *Rana sphenocephala*, *Pseudacris feriarum*, and *Pseudacris nigrita* mapped to 364 target loci in the *Xenopus tropicalis* genome (xenTro3). The 364 loci have been used broadly in vertebrate phylogenomic studies (e.g., Lemmon et al. 2012; Ruane et al. 2015; Prum et al. 2015; Barrow et al. 2018). After enrichment, 3-4 enrichment reactions were pooled in equal quantities for each sequencing lane and sequenced on two PE150 Illumina HiSeq 2500 lanes at the Translational Science Laboratory in the College of Medicine at Florida State University.

### Anchored Hybrid Enrichment Bioinformatics

The raw sequencing reads were processed following methods previously described (Lemmon et al. 2012; Rokyta et al. 2012; Prum et al. 2015; Ruane et al. 2015; Hamilton et al. 2016). Reads were demultiplexed with no mismatches tolerated and pairs were merged as in Rokyta et al. (2012). Merged reads were assembled using a quasi-denovo assembly approach described in Prum (2015) and Hamilton et al. (2015) using reference sequences for *Xenopus*, *Rana*, and *Pseudacris*. Alleles were phased using assembled read overlap information in a Bayesian statistical framework and orthology was assessed using sequence similarity. Extended bioinformatic methods are available in the Supplemental Methods.

### Polyploid Data Processing

To create species trees for polyploid complexes when the mode of polyploid formation is allopolyploid or otherwise unknown, whether using quartet-based or concatenated methods, it is first necessary to assign alleles to individual subgenomes. To begin this subgenomic assignment, alleles were phased using the allele phaser described in Pyron et al. (Pyron et al. 2016, but see Supplemental Methods). Initially, all samples were phased for four alleles, and specimen ploidy validation was conducted using our R package PloidyPal (www.github.com/wbooker/PloidyPal). The package PloidyPal was developed for the present study and uses pairwise genetic distances of the four phased alleles to determine the differential signal present from a known ploidy training sample. This signal is then used to predict ploidy for unknown individuals (full outlining of the algorithm can be found in the package README). To confirm our ploidy assessment, we also used the program nQuire (Weiß et al. 2018), which takes a different approach to ploidy estimation utilizing the distribution of raw sequences mapped to reference sequences. Samples identified as diploid from our assessment were then re-phased as diploid for subsequent analyses. Paralogs present in diploids that were identified during orthology assessment (see Anchored Hybrid Enrichment Bioinformatics Extended; Supplemental Methods) were then removed from any downstream analyses to prevent spurious assignment of diploid-tetraploid orthologs.

Similar to St. Onge et al. (2012), haplotypes of *H. versicolor* were then assigned to one of two putative subgenomes for phylogenetic analysis. Because we know *H. chrysoscelis* is at least one of the progenitors of *H. versicolor*, we calculated pairwise genetic distances of all *H. versicolor* haplotypes to the consensus *H. chrysoscelis* sequence and assigned the haplotype with the smallest distance to the VERS MIN subgenome (*H. chrysoscelis* subgenome) and the greatest distance haplotype to the VERS MAX subgenome (putative other subgenome). The intermediate two haplotypes in tetraploids were discarded for phylogenetic analyses. We realize this MIN/MAX characterization might predispose the phylogenetic analyses against autopolyploidy, but a variety of other methods we employed are not subject to this bias or usage of MIN/MAX delimitation and will help us understand the effect and limitations of this approach.

### Mitochondrial Genome Assembly

Although we enriched for nuclear loci only, non-target mitochondrial (mtDNA) reads are normally very high in copy number relative to the nuclear genome and so were also sequenced as bycatch, allowing us to extract mtDNA genomes from the enriched sequence data. We used SeqMan NGEN v. 13.0.1 (DNASTAR, Inc., Madison, WI) to map mtDNA reads to reference sequences. To develop reference sequences, we started with closely related species using a minimum match percentage of 70%. We first assembled *H. chrysoscelis* and *H. andersonii* mtDNA genomes using a previously assembled *H. andersonii* mtDNA genome as a reference (Warwick 2016). The assembled *H. chrysoscelis* reference was then used to develop mtDNA genomes for *H. versicolor*, *H. avivoca, H. arenicolor,* and *H. femoralis*. After developing the reference sequence for each species, individuals were mapped to the appropriate reference using a minimum match percentage of 85%.

### Nuclear Phylogenetic Analyses

To reconstruct the nuclear evolutionary history of the gray treefrog complex, we estimated phylogenies based on several different data sets. A total of 244 loci were used for phylogenetic analyses after removing sequences with evidence of paralogy in *H. chrysoscelis*. Phylogenetic analyses were conducted using RAxML under a GTR + G model (Lanave et al. 1984; Tavaŕe 1986; Lewis 2001) and for all analyses both gene and concatenated supermatrix trees were estimated. In the supermatrix analysis, the substitution model was partitioned by locus. We used the full dataset along with various subsets of the data to fully characterize the evolutionary history of the system. These data sets included: (1) full data set, (2) *H. chrysoscelis* and outgroups only, (3) VERS MIN and outgroups only, (4) VERS MAX and outgroups only, (5) VERS MIN plus *H. chrysoscelis*, and (6) VERS MAX plus *H. chrysoscelis*.

### Population Genetic Analyses

To characterize the population structure of the gray treefrog complex, we conducted STRUCTURE (version 2.3.4) (Pritchard et al. 2000; Falush et al. 2003) analyses to understand the population structure of *H. versicolor* and *H. chrysoscelis*. We conducted analyses with *H. chrysoscelis, H. versicolor* and with a dataset that included the complex sister species *H. avivoca.* We included all four SNP genotypes in *H. versicolor* and two SNP genotypes for each polymorphic site in diploids—coding the remaining two SNPs as missing data. Importantly, SNP identification is done prior to phasing and are not restricted to any subgenome. Each analysis was conducted both with 8,683 SNPs from 244 loci as well as 1 SNP per locus (244 total) with 10 replications for each K value from 2 to 7. MCMC chains were run for 150,000 samples with a 50,000 sample burn-in period and verified for consistency across each replicate. Analyses were input into STRUCTURE HARVESTER (version 0.6.94) (Earl and vonHoldt 2012) and CLUMPP (version 1.1.2) (Evanno et al. 2005; Jakobsson and Rosenberg 2007) to summarize across all runs. Final STRUCTURE plots were created with distruct (version 1.1) (Rosenberg and Nordborg 2002). We did not determine the optimal K value based on any one method—instead investigating all K values analyzed, because results derived from suboptimal K values can be highly informative and choosing a single K value by any method can be misleading (Pritchard et al. 2000; Evanno et al. 2005; Meirmans 2015). In addition to STRUCTURE analyses, we also conducted a principle components analysis (PCA) as implemented in the R package ‘hierfstat’ using the 8,683 SNPs previously generated. Finally, we estimated the distribution of polymorphisms across species and lineages as an assessment of incomplete lineage sorting (ILS). Specifically, we looked at the number of SNPs with fixed differences between *H. versicolor* and *H. chrysoscelis*, polymorphisms shared between the two species, and polymorphisms unique to each species. In addition to comparing polymorphisms across species, we also compared the number of polymorphisms for each category between *H. chrysoscelis* and each *H. versicolor* mitochondrial lineage individually (see Mitochondrial Phylogenetic Relationships and Coalescent Timing Results).

### Mitochondrial Phylogenetic Analysis and Coalescent Timing Estimation

To reconstruct the history of mitochondrial evolution in the gray treefrog complex, we used a published molecular clock estimate of a gene sequenced in this study to infer coalescent times and the potential dates of any whole genome duplication events. We conducted analyses in BEAST 2.5.0 (Drummond and Rambaut 2007; Bouckaert et al. 2014) to estimate divergence times of *H. versicolor* progenitors from *H. chrysoscelis* and the approximate time of clade divergences within each species. We used a random local clock model with a coalescent constant population tree prior. Due to the paucity of available fossils for calibration at the scale of our study and the lack of genes with direct clock rate measurements, we generated a distribution for the molecular clock rate based on published results from related taxa of the ND2 gene—defined with a gamma prior bounded from 0.005 to 0.015 with *α* = 0.001 and *β* = 5000 to account for variability in the clock rate found across studies (Macey et al. 1998; Crawford 2003a; Barrow et al. 2017). We ran the BEAST analysis with a single partition under a GTR substitution model and a gamma distribution for among-site rate variation (Lanave et al. 1984; Tavaŕe 1986; Lewis 2001). Finally, we conducted all analyses with a random initial starting tree for 10 million MCMC generations with 1 million generations for burn in.

### Migration and Descendance Model Testing

To determine the identity and number of ancestors that gave rise to tetraploid *H. versicolor*, we used the program migrate-n (version 4.2.14) (Beerli 2006; Beerli and Palczewski 2010) to test if *H. versicolor* originated a single or multiple times and to identify who the progenitors of *H. versicolor* were. More specifically, we investigated the probability of descendance with and without migration of each *H. versicolor* mitochondrial lineage from each *H. chrysoscelis* nuclear genetic lineage (as determined by STRUCTURE) and the other *H. versicolor* mitochondrial lineages. Due to computational constraints, we subsampled 50 random loci from the five individuals from each lineage that best encompassed the entire geographic and genetic diversity of that lineage (bolded individuals in Supp. Table 1, lineage in parentheses). We restricted the analyses to the MIN alleles in order to directly test whether these putative *H. chrysoscelis* alleles (present in the tetraploid *H. versicolor*) were in fact directly descended from *H. chrysoscelis* or instead descended from another *H. versicolor* lineage. We recognize that using MIN alleles alone may presuppose the selection of models including migration with *H. chrysoscelis*, but several other aspects of our study assess gene flow with *H. chrysoscelis* and any conclusions are not drawn from this analysis alone.

For each analysis, we assumed an HKY mutation model with a uniform prior of mutation scaled population sizes (*θ*) between 0 and 0.05 for each population and a uniform migration rate prior distribution between 0 and 1500. Prior distributions were chosen based on results from shorter preliminary analyses to ensure posterior distributions could be accurately estimated. We summarized each analysis across fifty replicates with four heated chains (chain temperatures: 1.0, 1.5, 3.0, 10^6^), and each replicate consisted of 4×10^7^ MCMC steps for each locus of which the first 1×10^7^ steps were discarded as burn-in. Finally, we used the Bezier approximation of log-marginal likelihoods calculated from each analysis to assess the probability of each model analyzed.

We conducted Migrate-n analyses using two steps. The objective of the first step was to determine the number of *H. versicolor* origins, such that if *H. versicolor* had a single origin, we would expect to observe only one *H. versicolor* lineage with a high probability of descendance from a *H. chrysoscelis* lineage. Conversely, if *H. versicolor* originated multiple times, we would expect to observe multiple *H. versicolor* lineages with a high probability of descendance from *H. chrysoscelis* lineages. To conduct this analysis, we first tested a set of divergence and migration models for each *H. versicolor* lineage individually, only altering which population was providing migrants into the chosen lineage for migration. To test descendance, we altered which *H. versicolor* or *H. chrysoscelis* population gave rise to the chosen lineage, and we allowed for migration if those two populations had any area of sympatry. The migration model was constrained to allow only migration between lineages in contact, with migration across ploidies restricted to diploid into tetraploid populations. We chose to allow for only unidirectional migration from diploid to tetraploids as polyploids are generally more tolerant of hybridization especially when involving differing ploidy levels and diploid to tetraploid gene flow is most commonly observed in nature (results from our ABC model testing and STRUCTURE analyses also support this limitation) (Stebbins 1947, 1971; Petit et al. 1999; Bogart and Bi 2013).

The objective of the second step was to test specific hypotheses about the evolutionary history of all *H. versicolor* lineages together. After running all analyses in the first step, we chose a final set of six different models to test different hypotheses of the history and formation of the complex. The final models chosen for this analysis were included based on the initial individual lineage analyses described above, but when two lineages had high support for descendance from the same population, we tested whether or not those lineages were independently formed from the source population or if they were formed through a stepping-stone migration pattern (Fig. 1; model 1-5). Additionally, we also tested these models against an independent polyploid formation model as suggested by Holloway et al. (2006) and Ptacek et al. (1994) (Fig. 1; model 6).

**Figure 1.**
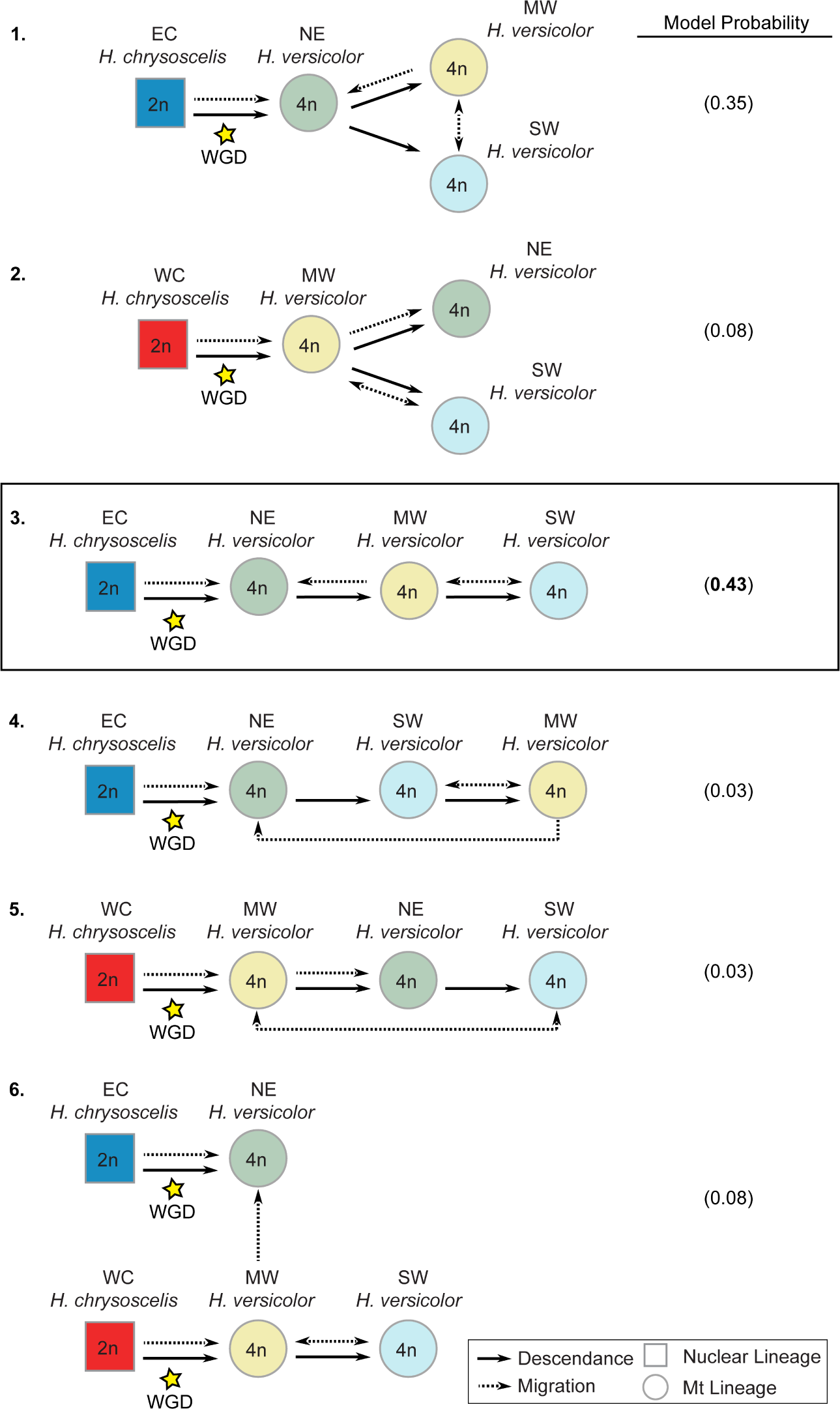
Models of *H. versicolor* descendance and migration used for the final migrate-n analysis and their model probabilities calculated using Bayes Factors from their Bezier approximation score. Boxed model demonstrates the model with the highest probability.

Although our estimate of the inheritance mode suggested the majority of loci are inherited tetrasomically (Fig. 3b), previous work has demonstrated inheritance polymorphisms in *H. versicolor* (Danzmann and Bogart 1982, 1983), we restricted migrate-n analyses to MIN alleles to ensure we were only testing whether a proportion of the *H. versicolor* genome originally descended from *H. chrysoscelis* or *H. versicolor* lineages and not whether the whole genome descended from a single lineage due to the probable contributions of extinct lineages to *H. versicolor* (see Mitochondrial Phylogenetic Analysis and Coalescent Timing Estimation Results). Though we were unable to directly test whether *H. versicolor* MIN alleles descended from extinct or extant *H. chrysoscelis* lineages here, the present analysis should identify which extant *H. chrysoscelis* lineage most similar to the extinct lineage contributed, if any. Indeed, results from individual lineage analyses support this assertion (i.e. detecting NE *H. versicolor* descendance from EC *H. chrysoscelis*, Table 2). A test of the possible contribution of extant and extinct *H. chrysoscelis* lineages to the original formation of *H. versicolor* is shown in the Approximate Bayesian Computation and Polyploid Speciation Model Testing Methods and Results.

**Table 2.** Bezier approximations of the marginal likelihood and the model probability from migrate-n analyses on descendance and migration likelihood for each *H. versicolor* mitochondrial lineage from each *H. chrysoscelis* nuclear genetic lineage or *H. versicolor* mitochondrial lineage. Bolded numbers represent the best supported model probability.

| Marginal Likelihood |  |  | Model Probability |  |  |  |
| --- | --- | --- | --- | --- | --- | --- |
| Descendance Model | NE | MW | SW | NE | MW | SW |
| EC No Migration | -117680.5 | -117668.1 | -117668.1 | 0.00 | 0.01 | 0.01 |
| EC Migration | -117664.6 | NA | NA | **0.47** | NA | NA |
| CC No Migration | -117697.1 | -117684.4 | -117684.1 | 0.00 | 0.00 | 0.00 |
| CC Migration | -117665.7 | -117667.5 | -117666.2 | 0.14 | 0.01 | 0.07 |
| WC No Migration | -117667.9 | -117710.0 | -117696.6 | 0.02 | 0.00 | 0.00 |
| WC Migration | NA | -117666.6 | -117666.2 | NA | 0.03 | 0.06 |
| NE No Migration | NA | -117663.2 | -117663.7 | NA | **0.88** | **0.81** |
| NE Migration | NA | -117666.4 | NA | NA | 0.04 | NA |
| MW No Migration | -117687.2 | NA | -117671.8 | 0.00 | NA | 0.00 |
| MW Migration | -117665.2 | NA | -117666.3 | 0.25 | NA | 0.06 |
| SW No Migration | -117665.9 | -117671.1 | NA | 0.12 | 0.00 | NA |
| SW Migration | NA | -117666.4 | NA | NA | 0.04 | NA |

### Approximate Bayesian Computation and Polyploid Speciation Model Testing

To ascertain the mode of polyploidization in the gray treefrogs, we used a modification of the framework outlined in Roux and Pannell (2015) in order to compare different models of polyploid evolution using Approximate Bayesian Computation (ABC). In brief, to conduct this analysis we: (1) sub-selected loci and individuals and calculated summary statistic means and standard deviations to generate our observed data, (2) estimated a nuclear molecular clock rate to be used for sequence simulation, (3) simulated multi-locus genetic data from biologically realistic prior parameter distributions for a selection of polyploid speciation models, (4) calculated summary statistic means and standard deviations for each simulation, (5) estimated model probability with ABC, and (6) estimated the posterior distributions of the parameters for the best supported models. These methods are explained in greater detail in the Supplemental Methods.

#### Generating Observed Data

We randomly selected 50 loci to use as the basis for our simulations and for calculating our observed summary statistics. Results from (2015) suggest that 20 loci are sufficient for distinguishing among polyploid speciation models, but their study only considers models without migration. We chose to use 50 loci to increase our power to correctly identify the best supported model when including more complex migration histories. We subsampled to 50 loci to accommodate computational limitations, and because randomization tests of the observed 50 locus estimate against a null of 1000 randomly generated 50 loci estimates demonstrates no test statistic calculated for 50 loci is outside of the null expectation (Supp. Table 2).

Alignments were first filtered to remove any columns with missing data to match the output of simulated sequences and accurately calculate the summary statistics for the observed data (Full list of summary statistics and values for the observed data, Supp. Table 2; but also see Use of the D3 Statistic for Polyploid Inference in the Supplemental Methods). We used alignments that included a random haplotype from each diploid *H. chrysoscelis* and the MIN and MAX alleles from *H. versicolor*. The ABC approach is best suited to using these data generated from our MIN/MAX separation, as the simulated evolutionary histories are conducted where the subgenomes are treated as their own populations, but the final summary statistics are calculated with the subgenomes treated as a single population (i.e. blind to any demarcation). Thus, our MIN and MAX assignment of observed haplotypes is most similar to the simulated data; however, because our summary statistic calculations do not assume any subgenomic assignment, this analysis is not biased towards any one model (from the model in Fig. 2a, MIN and MAX correspond to subgenomes A and B respectively).

**Figure 2.**
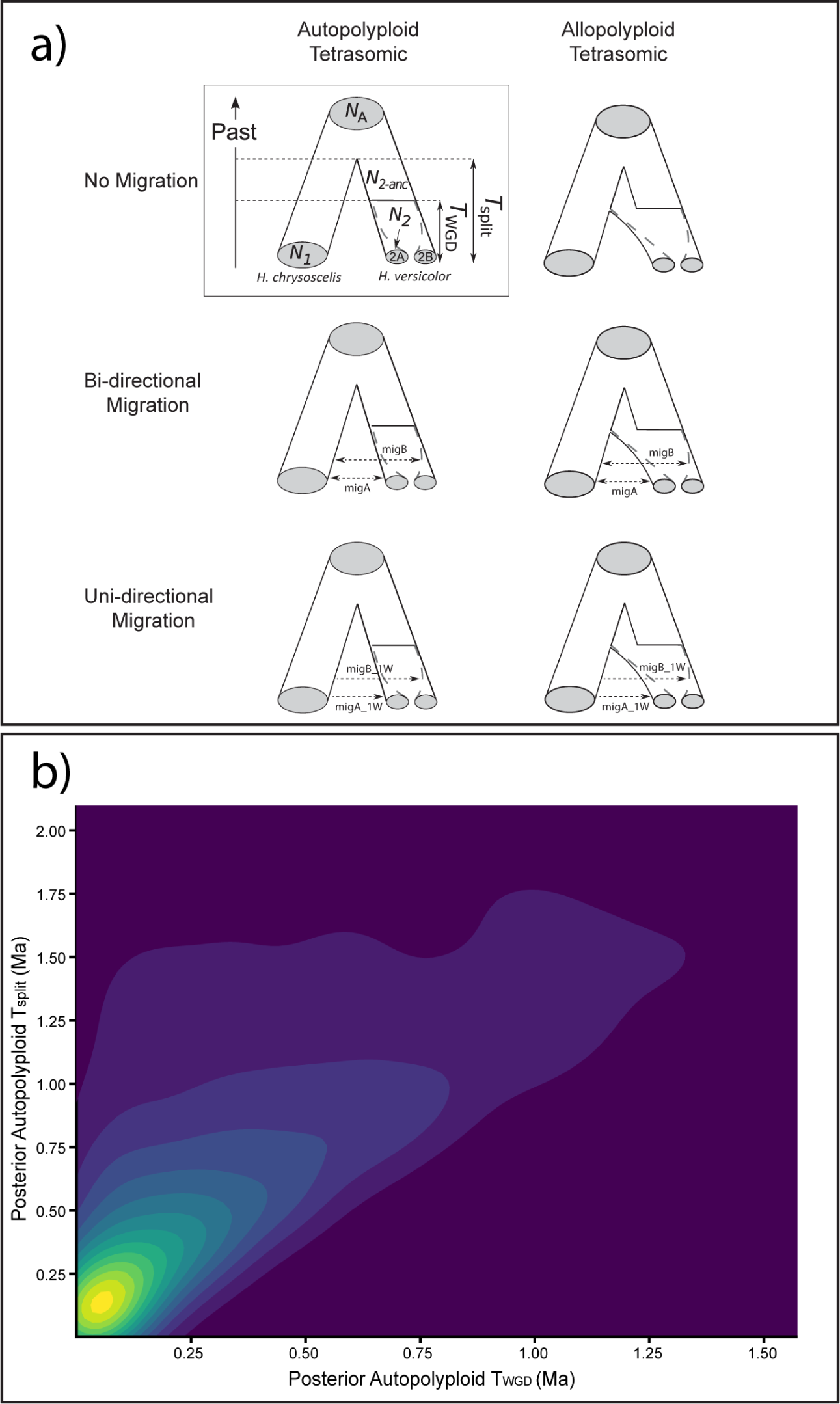
(a) The six polyploid speciation models and their parameters used in the ABC analysis (modified from Roux and Pannel 2015). The model inside the square depicts the parameters used for creating each model. (b) Two-dimensional density plot of posterior distributions for Tsplit and Twgd of the Autopolyploid Tetrasomic One-way Migration model used in the ABC analysis. Times are in millions of years ago (Ma).

**Figure 3.**
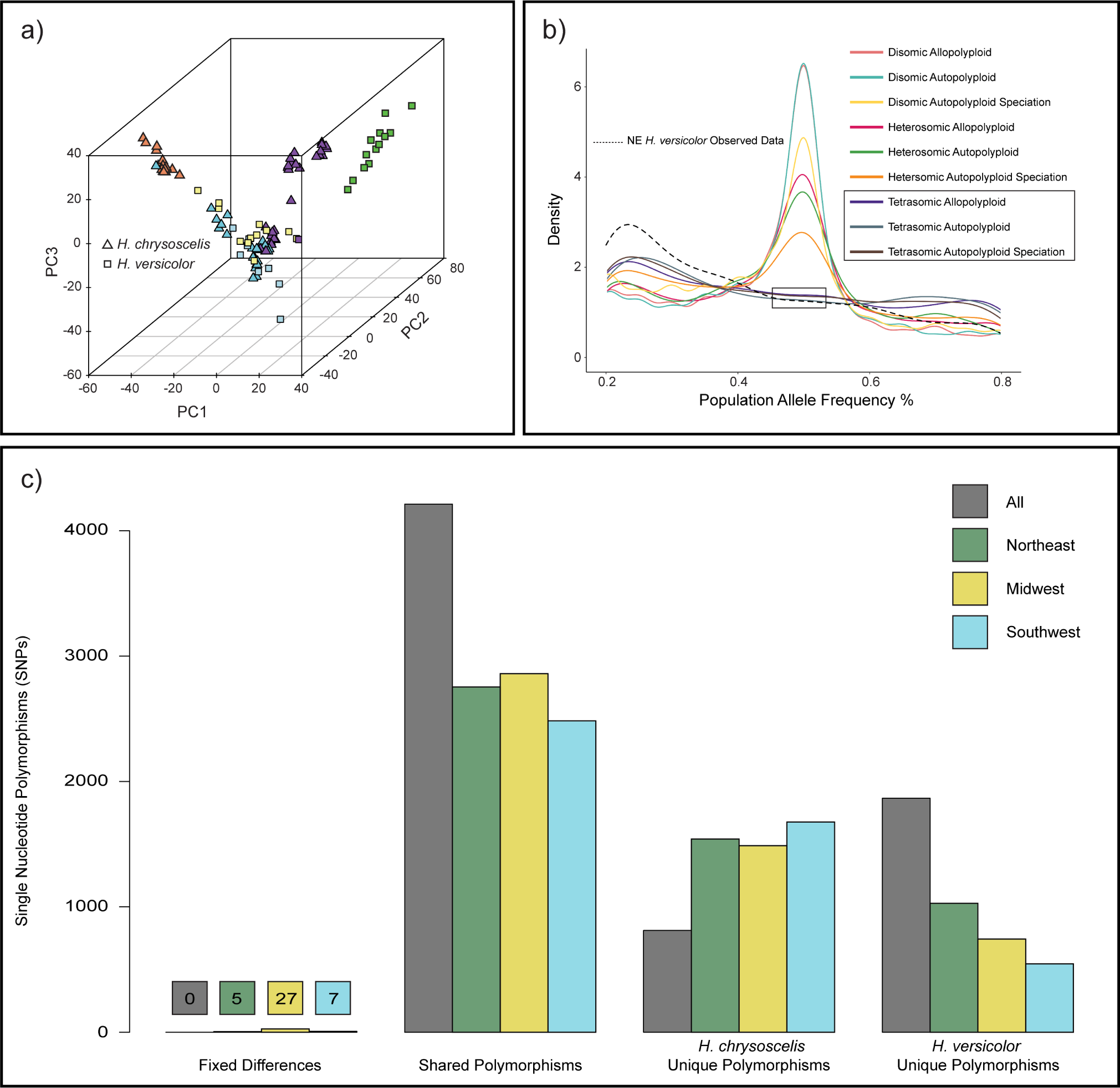
(a) Genetic PCA results for 8,683 SNPs across 244 loci. Squares and triangles represent *H. versicolor* and *H. chrysoscelis*, respectively. Fill color represents mitochondrial lineage and matches those presented in Fig. 5a-c colors. (b) Simulated and observed allele frequencies comparing nine speciation and inheritance mode combinations (one-way migration) to the observed allele frequencies of the NE *H. versicolor* lineage (shown as the dashed line). Simulations were done under a unidirectional migration model from the diploid to both A and B polyploid subgenomes. All heterosomic and disomic inheritance simulations produced allele frequencies with a significant number of alleles at 50% frequency, and tetrasomic inheritance simulations under all three polyploid speciation models did not produce a significant number of alleles at 50% frequency and were closest to the observed data. (c) Single nucleotide polymorphisms (SNPs) with fixed differences between *H. chrysoscelis* and *H. versicolor*, polymorphisms shared between *H. chrysoscelis* and *H. versicolor*, polymorphisms unique to *H. chrysoscelis*, and polymorphisms unique to *H. versicolor*. Comparisons were conducted between all *H. chrysoscelis* and *H. versicolor*, as well as between all *H. chrysoscelis* and each *H. versicolor* mitochondrial lineage.

#### Molecular Clock Rate Estimation

In practice, simulating the data for our models requires knowledge of the mutation rate for the chosen loci. Because we do not have molecular clock-rate estimates for our targeted loci, we used the software BEAST (version 2.5.0) (Drummond and Rambaut 2007; Bouckaert et al. 2014) to estimate a clock rate distribution for AHE loci (Barrow et al. 2018; Heinicke et al. 2018) in this group. We randomly selected 20 loci to calculate this distribution. For each locus, we set a gamma prior (*α* = 8.0, X = 0.88, offset = 2.1) on the coalescence time for the entire set of ingroup and outgroup samples based on the 95% credibility interval distribution of the coalescence time of that group as determined by our mitochondrial coalescent timing analysis. We used a GTR site model with a gamma distribution, a strict clock, and a Yule tree prior for each analysis. Each analysis was run for 1×10^7^ MCMC chains with a 2×10^6^ burn-in period. We used the mean value of the molecular clock rate across the 20 loci (multiplied by 10^-6^ to convert from mutations per site per million years to mutations per site per year) for our simulations, but allowed each simulated locus clock rate to be randomly chosen from a normal distribution with the mean being the mean clock rate across all loci (determined as 8.62×10^-10^ from our BEAST analysis, a rate similar to that found previously in frogs, e.g. (Crawford 2003b)) and with a standard deviation of 1×10^-10^ to allow for variability across loci.

#### Generating Simulated Data

We conducted 10^6^ multilocus simulations for six chosen models of polyploid evolution (possible model parameters Fig. 2a). Based upon results from our nuclear and mitochondrial phylogenetic analyses, we designed the simulated models to allow for either allopolyploidy (formation by hybridization of EC *H. chrysoscelis* and an extinct lineage) or autopolyploidy (whole genome duplication of an extinct lineage that split off from EC *H. chrysoscelis*) as the speciation mode. Chromosomal inheritance was limited to tetrasomic (all four tetraploid copies segregating freely) based on observed and simulated allele frequencies (see Site Frequency Spectrum Analysis methods; Fig. 3b); migration was either non-existent, asymmetric bi-directional to both tetraploid subgenomes, or unidirectional from the diploid to both subgenomes. Prior distributions for population size, Tsplit, TWGD and migration parameters were kept identical across all models. We used the Northeast *H. versicolor* mitochondrial lineage and the Eastern *H. chrysoscelis* nuclear genetic lineage to generate our observed data and the parameters for simulating each model based on results from our migrate-n analyses. We only included individuals whose genome contained only a small admixture portion from neighboring conspecific populations as determined from our STRUCTURE analysis. We chose this approach to ensure our observed data were most similar to the simulated data, which do not account for migration between populations other than the two used for this analysis.

To generate our priors, we used a modified version of the prior distribution script from Roux and Pannel (2015). To simulate sequences, we employed program msnsam (Hudson 2002; Ross-Ibarra et al. 2008). To calculate the summary statistics for each simulated dataset we used a modified version of mscalc (Ross-Ibarra et al. 2009; Roux et al. 2011, 2014; Roux and Pannell 2015). All scripts and complete instructions for using the pipeline are available in the Dryad repository or can be downloaded from www.github.com/wbooker/GTF_Polyploid_ABC.

#### Estimating Model Probability and Posterior Distributions

We used the R package ‘abc’ to estimate the probability of each model given our observed data (Csilĺery et al. 2012). We conducted each model probability estimation using a feed-forward neural network by nonlinear multivariate regression where the model itself is considered as an additional parameter to be inferred (scripts were modified from Leroy et al. (2017)). We selected 0.5% of the replicate simulations closest to the observed values of the summary statistics with a weighted Epanechnikov kernel. Computations were performed with 35 neural networks and 10 hidden networks in the regression. We then estimated the posterior distribution of the parameters for the three models with the highest probabilities with a neural network by nonlinear multivariate regression. For each model estimated, we conducted a logit transformation of the parameters for the 1,000 replicate simulations closest to the observed data (1.0% of the total number of simulations), and the posterior distribution of the parameters was then estimated using 35 trained and 10 hidden neural networks.

#### Assessment of Model Selection Robustness

To determine if the results from our ABC analysis provided more or less support for the chosen model, we assessed the robustness of ABC to accurately identify the correct model for a given model probability. To assess robustness, we simulated 1000 pseudo-observed datasets under both autopolyploid and allopolyploid models with tetrasomic and one-way AB migration (The two overwhelmingly supported models) and conducted ABC analyses for each of the pseudo-observed datasets. Robustness was assessed as 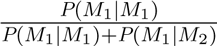.

### Site Frequency Spectrum Analysis

To better ascertain the mode of polyploid formation and chromosomal inheritance patterns, we estimated the distribution of allele frequencies, or the site frequency spectrum, for each *H. versicolor* lineage and compared these distributions to distributions generated from simulated data. The site frequency spectrum can be informative when analyzing genomic data and has been widely used to estimate demography and evolutionary histories (Gutenkunst et al. 2009). Because of the lack of recombination between subgenomes in the case of disomic inheritance (tetraploid subgenomes remain distinct, with segregation only between distinct subgenomic copies) and to a lesser extent heterosomic inheritance (mixed inheritance, with segregation between all four copies of some chromosomes, but only subgenomic copies for other chromosomes), the site frequency spectrum of a disomic or heterosomic polyploid should have an overabundance of alleles with a 50% frequency in the population as compared to tetrasomic polyploids (Hollister et al. 2012; Arnold et al. 2015). However, while one might observe such a pattern in an isolated population, it is unclear how various migration histories between diploids and tetraploids might affect our ability to discern a polyploid’s chromosomal inheritance pattern.

We simulated sequences and recorded allele frequencies using the same scripts, input files, and prior values as our ABC analysis for 45 polyploid speciation models (All combinations of inheritance, speciation, and migration patterns; outlined in Polyploid Speciation Model Testing Parameters in the Supplemental Methods). As in the ABC analysis, we assigned individuals to a genomic population only if their genome contained the few alleles from neighboring conspecific populations as determined from our STRUCTURE analysis. We ran 50,000 50-locus simulations for each model, using the same prior distributions of the parameters as our ABC analysis.

## Results

### Sequencing Summary

We successfully sequenced the target loci from all capture pools with minimal individual failures, producing a final dataset with 117 individuals (35 *H.* versicolor, 71 *H. chrysoscelis*, 7 *H. avivoca*, and the three outgroup taxa). Across all individuals, the average number of raw reads sequenced per individual was 7,723,547 (range: 2,490,670–25,206,362) and the average number of reads per locus was 1,984 (range: 16–6,039). After orthology assessment and manual trimming, the total number of locus alignments was 385, and after removing paralogs identified in *H. chrysoscelis*, we had a final number of 244 alignments. The average locus lengths for the 244 loci used were 1,380–1,383 sites depending on the dataset. The amount of missing data for nuclear datasets ranged from 2.6% to 2.8%.

The concatenated nuclear datasets consisted of 374,071-374,891 sites depending on the dataset, which included between 10,568 (2.83%) and 14,958 (3.99%) informative sites (Table 1). SNP extraction recovered 8,683 SNPs that were each flanked by 5 monomorphic sites with no missing data across *H. chrysoscelis* and *H. versicolor*. Whole mitochondrial genomes were recovered from all individuals sampled. The total number of sites recovered for the mitochondrial genome was 15,834, with 2,715 informative sites and 2.6% missing data.

### Polyploid Data Processing

Both ploidy assessment methods came to the same conclusions for each sample, and misidentified specimens were relabeled if an investigation of the field notes for that sample could confirm our results (i.e correctly identified in the field but misidentified or mislabeled afterward). Unconfirmed samples were removed from further analyses. Pairwise genetic distances between the average *H. chrysoscelis* and each sample in this study (separated into the MIN/MAX subgenomes for *H. versicolor*) demonstrate that on average *H. versicolor* is genetically very similar to *H. chrysoscelis*, with individual MAX subgenomes not exceeding a distance of 0.5% (Supp. Fig. 1). In comparison, *H. avivoca*, the sister species to the complex has an average distance of 1.23% to *H. chrysoscelis.* These results suggest that if *H. versicolor* were an allopolyploid derived from *H. chrysoscelis* hybridizing with an extinct species, at the very least the extinct species would have to have been a closer relative to *H. chrysoscelis* than *H. avivoca*, or that heterosomic inheritance in *H. versicolor* and/or hybridization between *H. versicolor* and *H. chrysoscelis* has eroded the genomic signal of a more distant relationship. Additionally, although we do see a difference between the putative MIN/MAX subgenomes in *H. versicolor* (average MAX=0.40%, MIN=0.27%), that difference is relatively small and less than the average distance of any individual *H. chrysoscelis* to the average *H. chrysoscelis* across all individuals (0.24%). Finally, measurements of within subgenome diversity demonstrate a greater diversity in the MAX subgenome, likely because restricting the MIN subgenome to the minimum difference from the average *H. chrysoscelis* restricts sequences to similarity in a single radial direction in sequence similarity/dissimilarity space, whereas the MAX subgenome sequences can be dissimilar in any radial direction (Supp. Table 3).

**Table 3.** Model probabilities from our ABC analysis under different polyploidization mode and migration histories. Probabilities are summarized across 20 independent runs. AB distinction for some models refers to migration with the diploid and both tetraploid subgenomes.

| Formation | Inheritance | Migration History | Model Probability |
| --- | --- | --- | --- |
| Autopolyploid | Tetrasomic | No Migration | 0.0002 |
| Allopolyploid | Tetrasomic | No Migration | 0.0002 |
| Allopolyploid | Tetrasomic | Bi-directional AB | 0.0124 |
| Autopolyploid | Tetrasomic | Bi-directional AB | 0.0250 |
| Allopolyploid | Tetrasomic | Uni-directional AB | 0.3940 |
| Autopolyploid | Tetrasomic | Uni-directional AB | 0.5682 |

### Nuclear Phylogenetic Relationships

Phylogenetic analyses of nuclear data provided support that *H. versicolor* may harbor alleles from an unsampled, apparently extinct population or species (Fig. 4a,e-f, *H. versicolor* sequences outside of the *H. chrysoscelis* clade). In addition, these data showed evidence of clear genetic breaks across geography for both *H. chrysoscelis* and *H. versicolor*. Individual gene trees exhibited low bootstrap support, likely due to the low information content at each locus at this shallow phylogenetic scale. Concatenated analyses of all subset nuclear data sets, however, generally had high bootstrap support for informative branches. Conversely, low branch bootstrap support for some analyses may be informative as well for identifying a lack of variation between *H. versicolor* and *H. chrysoscelis* genes (e.g., Fig. 4e).

**Figure 4.**
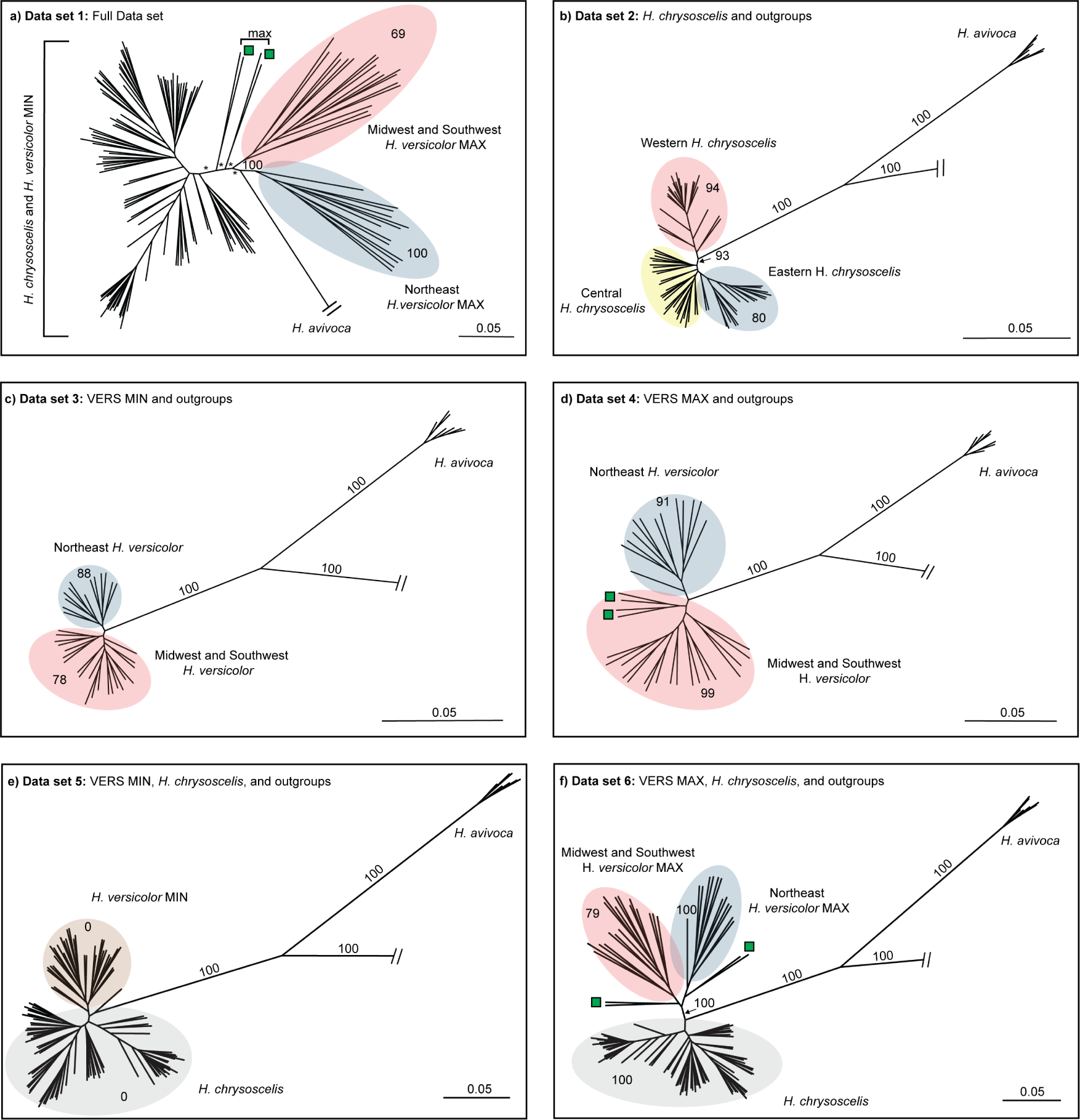
RAxML phylogenies for concatenated alignments across 244 AHE loci. Colored ellipses highlight clades of interest. Bootstrap values are only reported on branches informative for this study (full trees with tip labels and bootstrap support at all nodes can be found in the appendix). Stars represent bootstrap support *<*50. Green squares indicate clades that suggest potential past connectivity of *H. versicolor* NE and SW lineages (see Discussion). Scale bar and branch lengths represent substitutions per site. Outgroup relationships shown in Supp. Fig. 2.

The concatenated analysis using the full dataset (data set 1; Fig. 4a) recovered a topology similar to those of subset analyses (data sets 2-6; Fig. 4b-f), but ultimately had low bootstrap support values at most branches. The two subset analyses with both *H. versicolor* and *H. chrysoscelis* (Fig. 4e-f) resulted in topologies with high bootstrap support that provide evidence that *H. versicolor* has a large proportion of alleles that came from a population or species that was sister to all extant *H. chrysoscelis*. Additionally, within the two subset analyses including MIN or MAX *H. versicolor* and outgroups only, the recovered pattern suggests extant *H. versicolor* is separated into two general clades—one eastern nuclear genetic lineage consisting of all sampled individuals from WV, VA, MD, NJ, NY, CT, ME, and one western nuclear genetic lineage consisting of all other sampled *H. versicolor* (Fig. 4c-d). Consistent with previous work (Ralin and Selander 1979; Holloway et al. 2006), our analysis of *H. chrysoscelis* and outgroups identifies two clades within *H. chrysoscelis*, a Western clade and an Eastern + Central clade, with a monophyletic Eastern clade nested within the Central lineage (Fig. 4b; Fig. 5c).

**Figure 5.**
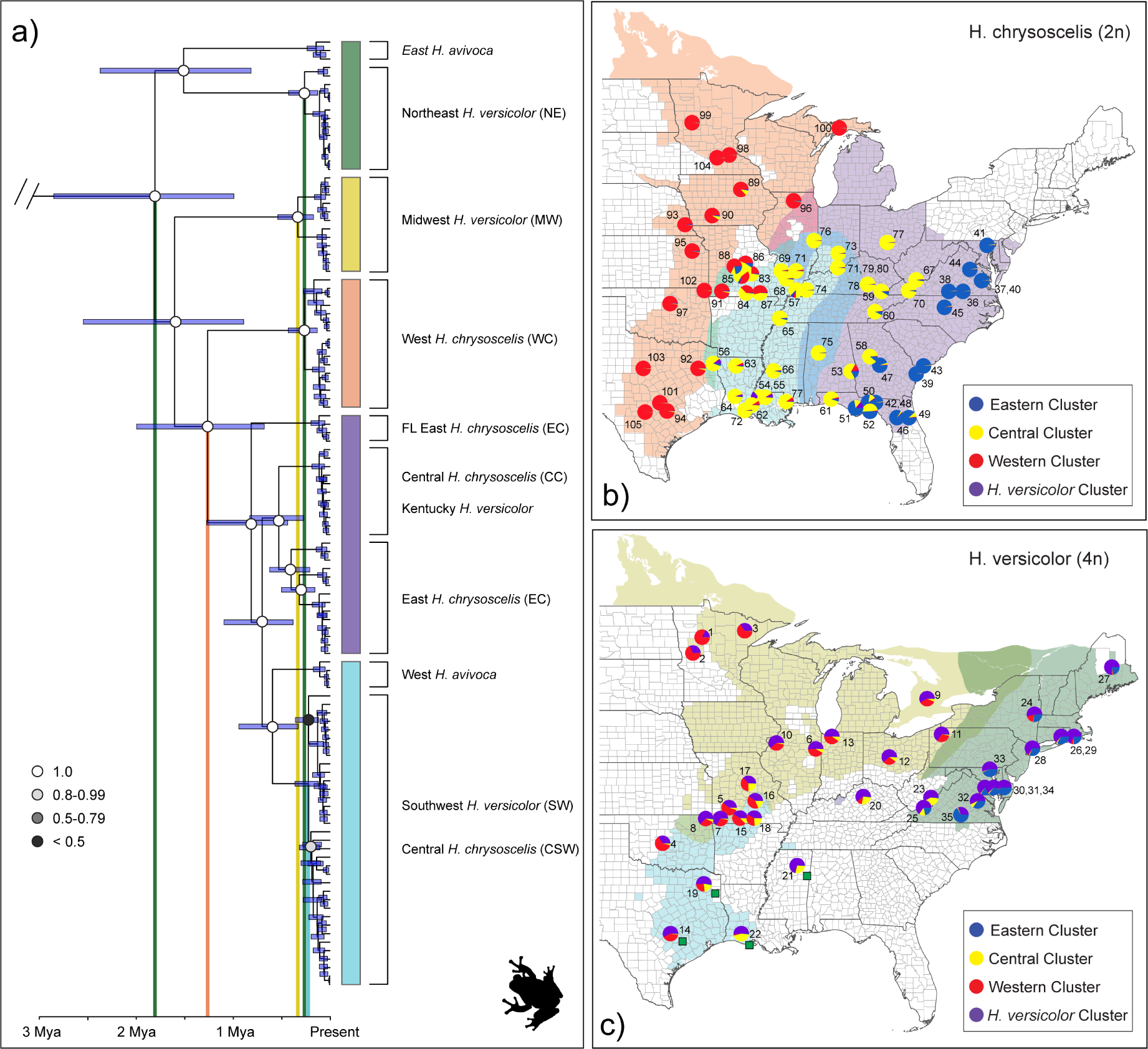
(a) Dated whole-genome mitochondrial phylogeny from BEAST analysis of all ingroup and outgroup taxa (*H. andersonii, H. arenicolor,* and *H. femoralis* not shown, see Supp. Fig. 4). Colored bars right of the phylogeny highlight mitochondrial clades. Circle color on nodes represent posterior values for those nodes and are only reported for branches informative for this study (Full tree with all tip and node labels is available in the Dryad repository, Fig. 3). From left to right, vertical bars show mean timing of coalescence for: 1) *H. avivoca*, *H. versicolor,* and *H. chrysoscelis*; 2) Eastern/Central *H. chrysoscelis* and Western *H. versicolor*, 3) all MW *H. versicolor*; 4) all NE *H. versicolor*; and 5) all SW *H. versicolor* and the Central *H. chrysoscelis* with which they share a monophyletic mitochondrial clade. (b) Distribution map of *Hyla chrysoscelis*. Background colors indicate putative ranges of mitochondrial lineages, circles represent nuclear STRUCTURE results for *H. versicolor* and *H. chrysoscelis* analysis at K=4 with one SNP per locus. K=4 is visualized here because this analysis most reflected the topology from our phylogenetic analysis (Fig. 4b), did not include additional clusters that were uninformative, and overall was most useful for visualization of the complex population structure. Numbers correspond to Map ID number in Table 1 and Fig. 5-6. Green squares next to Texas, Louisiana, and Tennessee samples correspond to individuals that had ambiguous relationships in RAxML analyses. (c) Distribution map of *Hyla versicolor*. Background colors indicate putative ranges of mitochondrial lineages, circles represent STRUCTURE results for *H. versicolor* and *H. chrysoscelis* analysis at K=4 with one SNP per locus. Background colors indicate putative ranges of mitochondrial lineages, circles represent nuclear STRUCTURE results for *H. versicolor* and *H. chrysoscelis* analysis at K=4 with one SNP per locus. Numbers correspond to Map ID number in Table 1 and Fig. 5-6.

Additionally, other patterns from our nuclear phylogenetic analysis suggest a more complex evolutionary history than can be explained by simple bifurcations. Green squares on the phylogenies in Fig. 4 and map in Fig. 5b indicate *H. versicolor* individuals from Texas, Louisiana, and southwest Tennessee that were consistently placed on the phylogeny nearby but outside the clades expected from their geographic location. These patterns may be evidence of hybridization between *H. chrysoscelis* and *H. versicolor* —a conclusion reached by several other analyses in this study.

Finally, the taxonomic relationships of the outgroup taxa from our nuclear analyses demonstrate a unique topology that conflicts with previous estimates (Fig. 2) (Duellman et al. 2016; Faivovich et al. 2005). For all concatenated analyses, we recovered species relationships that separate *H. versicolor* and *H. chrysoscelis* from *H. avivoca*, *H. andersonii*, *H. arenicolor*, and *H. femoralis* as two monophyletic clades with high bootstrap support. Within the outgroup clade, *H. avivoca* forms a monophyletic clade separate from the other outgroup taxa, and within *H. avivoca*, we recover a topology that separates East (AL, GA, TN) and West (MS, LA) *H. avivoca* with high support.

### Mitochondrial Phylogenetic Relationships and Coalescent Timing

Whole mitochondrial genome analyses recovered a topology similar to that found in the single-gene mitochondrial study of Ptacek et al. (1994), but our increased gene and taxon sampling improved phylogenetic estimates and clarified the origin of lineages (Table 1; Fig. 5a; Supp. Fig. 3). Five main clades were delineated: one containing *H. versicolor* only (Midwest, MW); one consisting of northeastern *H. versicolor* and eastern *H. avivoca* (Northeast, NE); one including western *H. avivoca,* southwestern *H. chrysoscelis* (CSW), and southwestern *H. versicolor* (Southwest, SW); a western *H. chrysoscelis* clade (West *chrysoscelis*, WC); and one paraphyletic group containing East *H. chrysoscelis* (EC), Central *H. chrysoscelis* (CC) and one *H. versicolor* from Meade County Kentucky (Fig. 5a). All major branches had posterior probabilities of 1, apart from internal nodes within the SW/CSW clade where *H. versicolor* and *H. chrysoscelis* are not individually monophyletic.

The whole mitochondrial genome analysis for our outgroup taxa recovered species level relationships with high posterior probability, but a topology that conflicts with the topology estimated by our nuclear analyses (Fig. 4). In the mitochondrial analysis, we recovered a topology that places the ingroup (including *H. avivoca*) as sister to *H. andersonii*, and this clade is sister to *H. arenicolor*. Finally, this analysis delineates *H. femoralis* as sister to all other species used in this study.

We recovered mitochondrial coalescent timing estimates with narrow 95% credibility intervals for all nodes with a mean clock rate of 0.906% per lineage per million years. Date estimates of importance can be found in Supp. Table 4, but briefly, these estimates suggest coalescent timing of all identified *H. versicolor* lineages as sometime within the last 430,000 years (NE: mean 0.262 Ma, 95% CI 0.125-0.426 Ma; MW: mean 0.338 Ma, 95% CI 0.131-0.430 Ma; SW: mean 0.223 Ma, 95% CI 0.120-0.360 Ma). This analysis placed the coalescent timing of all *H. chrysoscelis* within the last 1.99 Ma (mean 1.26 Ma, 95% CI 0.675-1.99 Ma).

### Population Genetic Analyses

Within *H. chrysoscelis*, our STRUCTURE analysis across multiple K values suggest there are three distinct clusters representing Western, Central, and Eastern *H. chrysoscelis* (Fig. 5c; Supp. Fig. 5; Supp. Fig. 6). This result is consistent with our nuclear phylogenetic and whole mitochondrial genome phylogenetic results (Fig. 4b; Fig. 5a), however the nuclear phylogeny places Eastern *H. chrysoscelis* nested within the Central lineage, and the mitochondrial phylogeny shows two divergent mitochondria segregating within the center of the Eastern/Central lineage’s range. Although individual *H. chrysoscelis* generally have a *>*90% identity match to a single cluster, some geographic locales contain several individuals with significant genetic contributions from neighboring or sympatric lineages, suggesting hybridization between *H. chrysoscelis* lineages in contact zones.

**Figure 6.**
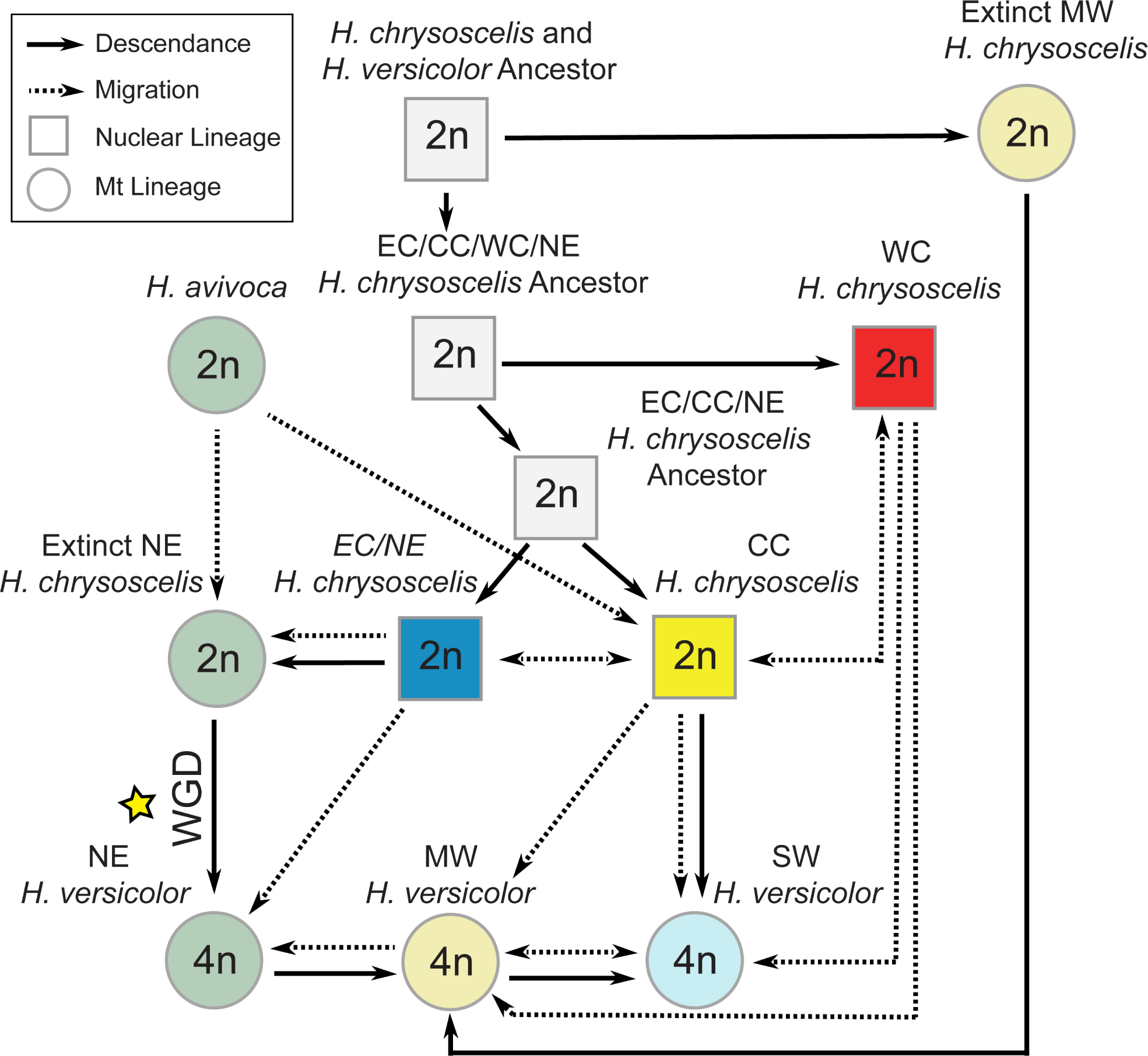
Network model of speciation and evolution proposed from the results of this study. Background colors of circles represent mitochondrial lineages apart from ancestral lineages which are colored white. Solid arrows point to descendants of a given lineage. Dashed lines demonstrate current migration for existing lineages or past migration for extinct lineages, with arrows indicating the direction of migration. Circles represent populations identified by their mitochondrial genome (*H. versicolor, H. avivoca*, and extinct lineages), squares represent populations defined by their nuclear genome (*H. chrysoscelis*). The yellow star indicates the proposed single whole genome duplication event that led to the formation of *H. versicolor*.

Within *H. versicolor*, all individuals show evidence of a unique genetic influence that is generally absent from *H. chrysoscelis* outside of minor frequencies (shown in purple, Fig. 5b; Supp. Fig. 5; Supp. Fig. 6). This pattern is consistent across several K values in both the single SNP per locus and all SNPs analyses. Interestingly, an additional analysis that included *H. avivoca*, showed evidence that this distinct *H. versicolor* cluster might have descended from *H. avivoca* (purple cluster at K=3-4, Supp. Fig. 7). This observation is further supported by the mitochondrial tree, which shows that eastern *H. avivoca* share a recent mitochondrial ancestor with Northeast *H. versicolor* (Fig. 5a). Additionally, some support was found at K=3 that alleles contributed to *H. versicolor* from *H. avivoca* were first integrated into the Eastern *H. chrysoscelis* genome. Although not conclusive, this pattern may suggest a single origin of *H. versicolor* from an ancestral population most closely related to Eastern *H. chrysoscelis*.

Additionally, we also found similar Western, Central, and Eastern clusters in *H. versicolor* as observed in *H. chrysoscelis*. Aside from the unique genetic influence in *H. versicolor* (purple cluster), other cluster proportions within individuals are remarkably similar to nearby *H. chrysoscelis* populations—especially so when *H. versicolor* and *H. chrysoscelis* are sympatric (Fig. 5c; Fig. 5b). We found a similar pattern in our genetic PCA analysis (Fig. 3a), with Midwest and Southwest *H. versicolor* being generally indistinguishable from Eastern and Central *H. chrysoscelis* across axes 1-3 (PCA axes 1-3 represent 4.8%, 3.2%, and 2.5% of the variability, respectively). Genetic PCA results also show Northeast *H. versicolor* as distinct from all other *H. versicolor* and *H. chrysoscelis* but most similar to Eastern *H. chrysoscelis*. The divergence of NE *H. versicolor* from MW and SW lineages is also seen at higher K values from our analysis that included *H. avivoca*(green cluster, Supp. Fig. 7.

Our assessment of the distribution of SNPs demonstrate significant ILS between *H. chrysoscelis* and *H. versicolor* 3c. We found no fixed differences across the two species as a whole, and very few fixed differences between *H. chrysoscelis* and each individual *H. versicolor* lineage. These results also demonstrate the majority (*>*4000) of SNPs we recovered represent shared polymorphisms between *H. chrysoscelis* and *H. versicolor*, indicating significant ILS in this complex. Finally, this assessment also shows greater nucleotide diversity within *H. versicolor* than in *H. chrysoscelis* when the two species are compared collectively, but the reverse relationship when comparing *H. chryoscelis* to each *H. versicolor* lineage individually. The lineage level analysis also demonstrates a difference in nuclear diversity across lineages that supports NE origin and stepping stone model identified from our Migration and Descendance Model Testing Results (model 3; Fig. 1).

### Migration and Descendance Model Testing

Our single population migrate analyses suggest that NE *H. versicolor* originated from Eastern *H. chrysoscelis*, and that MW and SW *H. versicolor* are descendants of NE *H. versicolor* —providing support that *H. versicolor* arose from a single whole genome duplication event (Table 2). Indeed, the Northeast *H. versicolor* lineage was the only lineage that had a significant probability of being a descendant of any one *H. chrysoscelis* lineage, with descendance from Eastern *H. chrysoscelis* having the highest probability (0.47 model probability, Table 2). For both the Midwest and Southwest *H. versicolor* lineages, the best supported model was one of descendance without current migration from the Northeast *H. versicolor* lineage (0.88 and 0.81 model probability, respectively). When we tested the final set of six models, however, we found higher support for a stepping-stone model, where MW *H. versicolor* descended from NE *H. versicolor,* and SW *H. versicolor* descended from MW *H. versicolor*. Importantly, the above model was not overwhelmingly supported (0.43 model probability; Fig. 1, model 3), and a model of independent origination of MW and SW *H. versicolor* from NE *H. versicolor* also received considerable support (0.35 model probability; Fig. 1, model 1). We found little support for any other models, including the model of multiple origins.

When considering migration patterns, almost all models tested in our initial analyses had a higher log-likelihood when allowing for migration between any two lineages when appropriate. Specifically, all models improved when we allowed for unidirectional migration of diploid lineages into sympatric tetraploid lineages, and nearly all models improved when we allowed for migration between neighboring tetraploid lineages. The only model comparison that had worse support when allowing for migration was our Midwest *H. versicolor* descending from Northeast *H. versicolor* comparison, in which a model that didn’t allow for migration from Northeast *H. versicolor* had higher support (Table 2).

### Approximate Bayesian Computation Polyploid Speciation Model Testing

The best supported model from our ABC analysis suggests *H. versicolor* originated as an autopolyploid from an extinct *H. chrysoscelis* lineage, and that there is unidirectional migration from Eastern *H. chrysoscelis* into Northeastern *H. versicolor* (model probability = 0.57); Table 3). In addition, almost all model support is concentrated across both unidirectional migration models, with the allopolyploid unidirectional migration model also receiving a high level of support (model probability = 0.39). A linear discriminate analysis (LDA) of each of the six models shows that although models with different migration parameters can generally be distinguished in LDA space, when there is migration between diploids and tetraploids, autopolyploid and allopolyploid models become almost indistinguishable (Supp. Fig. 8). Importantly, however, our observed data (star in Supp. Fig. 8) are well within LDA space and therefore the priors used for our simulations. Our robustness assessment shows that our ABC analysis is able to distinguish allopolyploid and autopolyploid models when there is migration, but this assessment does not provide any additional confidence that the chosen model is the true model outside of the originally estimated probability (Supp. Fig. 9).

Posterior distributions for the best supported model suggest that the extinct lineage that led to the autopolyploid formation of *H. versicolor* split from Eastern *H. chrysoscelis* around 2.88×10^5^ years ago (peak distribution value (PDV): 2.88×10^5^; 90% CI: 1.02×10^3^ - 3.22×10^6^), and that the whole genome duplication event occurred 1.00×10^5^ years ago (PDV: 1.00×10^5^; 90% CI: 2.4×10^1^ - 1.84×10^6^) (Fig. 2b; all population size and timing posteriors Supp. Fig. 11). Alternatively, under the second-best supported model, the peak posterior distribution value suggests much earlier Tsplit and Twgd times at 4.19×10^6^ (PDV: 4.19×10^6^; 90% CI: 2.00×10^6^ - 8.11×10^6^) and 2.96×10^6^ (PDV: 2.96×10^6^; 90% CI: 9.07×10^5^ - 2.96×10^6^) years ago respectively (Supp. Fig. 11). Though largely overlapping with the prior, the posterior values for the autopolyploid model are consistent with the coalescent times observed in our mitochondrial phylogenetic analysis (Supp. Table 4), while the posterior for Tsplit and Twgd times under an allopolyploid model would predate the coalescent times of all *H. chrysoscelis* and *H. versicolor* from the same analysis (Fig. 5a). Given that the nuclear clock rate we used to simulate the data was estimated given the coalescent times from our mitochondrial phylogenetic analysis, the dates estimated by the ABC analysis should be approximately within the estimated coalescent timing distribution, even if that estimate is incorrect. These results, in our view, add further support to an autopolyploid model, suggesting that for an allopolyploid model to be supported, the Tsplit and Twgd times would need to be well outside of what is biologically reasonable.

### Site Frequency Spectra of Simulated Polyploid Models

Site frequency spectra at intermediate allele frequencies were similar across all simulated migration histories—suggesting that significant deviations from tetrasomic inheritance should be detectable as an excess of observed alleles at 50% frequency in the population regardless of migration history, given some differentiation has occurred between subgenomes since WGD for autopolyploid species with disomic or heterosomic inheritance (Fig. 3b; Supp. Fig. 10). The observed frequencies for all mitochondrial *H. versicolor* lineages were nearly identical and most similar to those observed by the simulated tetrasomic models. Previous segregation studies have suggested that *H. versicolor* is not entirely tetrasomic, with some genes segregating in a disomic fashion (Danzmann and Bogart 1982, 1983). Importantly, the simulated heterosomic models are produced by randomly choosing some number of disomic loci from a uniform prior—meaning the density histogram of allele frequencies reflects a genome with ~50% disomic inheritance. Genomes with only one or a few chromosome sets forming bivalents may be indistinguishable from completely tetrasomic genomes without significant sampling across the genome. Thus, it is possible there is some level of disomic inheritance in the *H. versicolor* genome; however, the overwhelming pattern of chromosomal inheritance is tetrasomic, and we are confident in eliminating heterosomy and disomy from our chosen set of models.

## Discussion

As evidenced here, model-based approaches are an important tool for to testing alternative evolutionary hypotheses to disentangle the complex relationships in polyploid systems. Ours is the first study that has not relied solely on descriptive methods to delineate the history of gray treefrogs, and in its totality, the evidence presented above largely disagree with the previous conclusions from Holloway et al. (2006). Specifically, rather than supporting multiple allopolyploid origins, our study instead suggests that *H. versicolor* most likely formed via a single autopolyploid whole genome duplication event, and that current lineages of *H. versicolor* are a result of repeated backcrossing with extant and extinct lineages of *H. chrysoscelis* 6. Several lines of evidence from our study also corroborate previous observations that *H. versicolor* hybridizes with *H. chrysoscelis* when they are sympatric (Gerhardt et al. 1994) and the hypothesis that *H. chrysoscelis* alleles regularly introgress into the *H. versicolor* genome to create high levels of heterozygosity (Ralin and Selander 1979; Bogart and Bi 2013). Finally, multiple results from this study suggest that the sister species to the complex, *H. avivoca,* has played an ongoing role in the genetic history of this group as well.

### Mode of Polyploid Formation

Multiple lines of evidence suggest that *H. versicolor* is an autopolyploid that formed via a single whole genome duplication event from an extinct Northeast *H. chrysoscelis* lineage that was genetically most similar to present day Eastern *H. chrysoscelis*. Autopolyploidy is indicated both through our observation of intermediate allele frequencies that reflect simulated tetrasomic polyploid models, pairwise genetic distance of *H. versicolor* to *H. chrysoscelis*, as well as our results from ABC polyploid speciation model testing (Fig. 3b; Supp. Fig. 1; Table 3). Although an Allopolyploid model was also moderately supported from our ABC analysis, the posterior estimates of Tsplit and TWGD times under this model (Supp. Fig. **??**) fall well outside the times estimated by our mitochondrial coalescent analysis (Supp. Table 4) that were used to estimate the nuclear clock rate for the ABC analysis. Autopolyploidy is also most consistent with the phenotypic and ecological similarity of *H. versicolor* and *H. chrysoscelis*. The two species are indistinguishable apart from their calls and the associated female responses, and while there is some evidence to suggest the two species partition their calling sites when sympatric, no other study has been able to identify an ecological or phenotypic difference between *H. versicolor* and *H. chrysoscelis* that is not a direct consequence of polyploidy itself (e.g. increased cell size) (Ralin 1968; Kamel et al. 1985; Ptacek 1992).

The Holloway et al. (2006) study is the only study of which we are aware that makes a definitive conclusion of allopolyploidy in *H. versicolor*. Here, we draw similar conclusions to those outlined by Bogart and Bi (2013)—namely that the Holloway et al. (2006) observation of alleles in *H. versicolor* not present in extant *H. chrysoscelis* is also compatible with an autopolyploid origin of *H. versicolor*. This conclusion suggests rather that *H. chrysoscelis* has long been a widespread species across eastern North America, and some historical populations that contributed genes to *H. versicolor* are now extinct. However, an additional result from our study could also suggest allopolyploidy—the phylogenetic separation of MAX subgenomes from a MIN and *H. chrysoscelis* clade shown in Fig. 4a despite the majority of loci having tetrasomic inheritance. From the totality of evidence presented in this study, we do not believe this is due to allopolyploidy, but rather the high levels of ILS observed (Fig. 3c, the possibility that some loci are inherited disomically (e.g., Danzmann and Bogart 1982, 1983), recent gene flow from diploids into tetraploids 3, and the overall effect grouping alleles into two classes by divergence has on phylogenetic assessment. Finally, as Bogart and Bi (2013) and Bogart et al. (2020) suggest but was originally proposed by Bogart and Wasserman (1972), our results are not inconsistent with the hypothesis that *H. versicolor* formed via a triploid intermediary (i.e. triploid bridge). Given the highly reticulate nature of this complex, however, we are unable to provide any evidence to support or reject their hypothesis outside of our support for autopolyploidy.

### Identity of Ancestors and Number of Origins

Our results suggest that *H. versicolor* arose from a single whole genome duplication event of a now extinct lineage of *H. chrysoscelis* that was most genetically similar to present day Eastern *H. chrysoscelis*. A single origin is supported by our migrate-n model testing, with support for descendance from any *H. chrysoscelis* population only occurring for the Northeast *H. versicolor* model (Table 2). Similarly, our other single-lineage migrate-n model tests demonstrate high support that Midwest and Southwest *H. versicolor* are descendants of Northeast *H. versicolor* (Table 2). However, a final model test suggests that Southwest *H. versicolor* is not a direct descendant of Northeast *H. versicolor*, but rather a descendant of Midwest *H. versicolor* populations which had descended from Northeast *H. versicolor* (Fig. 1, model 3). This model is further supported from our ILS assessment, which demonstrates decreased nucleotide diversity in Midwest *H. versicolor* compared to Northeast *H. versicolor*, and decreased nucleotide diversity in Southwest *H. versicolor* compared to Midwest *H. versicolor*.

A single origin is also supported by the presence of a unique genetic cluster found in all *H. versicolor* from our STRUCTURE analyses that may be derived from *H. avivoca* through an intermediate Eastern *H. chrysoscelis* conduit (Supp. Fig. 7). Mitochondria from the Northeast *H. versicolor* clade are most closely related to eastern *H. avivoca* mitochondria (Fig. 5a), but the migrate-n model tests (Table 2), pairwise genetic distance of *H. versicolor* and *H. chrysoscelis* (Supp. Fig. 1), nuclear phylogenetic analysis (Fig. 1), and genetic PCA (Fig. 3a), all suggest Northeast *H. versicolor* descended from the much more recently derived Eastern *H. chrysoscelis* population. Incomplete lineage sorting (ILS) is not likely to produce this pattern, but hybridization can, if some Eastern *H. chrysoscelis* population hybridized with *H. avivoca* prior to the formation of *H. versicolor*. This conclusion would also explain why we see clustering of *H. avivoca* with Eastern *H. chrysoscelis* and Northeastern *H. versicolor* at K=3 (Supp. Fig. 7). The results reported here largely corroborate the Romano et al. study (1987) that suggested a single origin, but this study suggests an origination from Northeast *H. versicolor,* whereas their best estimate of the location of origin for *H. versicolor* was somewhere in the central part of the range (i.e., MW *H. versicolor*).

While the bulk of evidence outlined in this study best supports a single origin of *H. versicolor*, some aspects of this study instead could provide support for multiple autopolyploid origins of *H. versicolor*. First, our mitochondrial phylogenetic analyses confirm that in addition to an extinct population of Northeast *H. chrysoscelis*, there was at least one more population of *H. chrysoscelis* that is now extinct and whose genome is now at least partially retained in Midwest *H. versicolor* (Fig. 5a). Second, Midwest and Northeast *H. versicolor* are largely genetically distinct, and the coalescent times of Midwest and Northeast *H. versicolor* ancestors are nearly identical. However, the only direct test for the origins of polyploidy overwhelmingly support a single origin (Fig. 1; Table 2). While future, more sophisticated analyses may suggest different conclusions than our own, patterns alone can be misleading and should not be solely relied on for drawing conclusions in polyploid systems.

### Complex Population Origin, Composition, Timing, and Hybridization

The totality of evidence from this study suggests that the North American gray treefrogs are a highly reticulate group with current ongoing exchange of alleles across polyploid and diploid lineages. Indeed, only one line of evidence from this study does not support the conclusion that any two lineages will hybridize whenever they come into contact—regardless of ploidy (migrate-n analysis suggests NE *H. versicolor* is not contributing migrants to MW *H. versicolor*, Table 2). The strong evidence of *H. chrysoscelis* to *H. versicolor* gene flow from this study supports recent findings from Bogart et al. (2020) that demonstrated evidence of local gene flow between the two species, but our work expands this observation to demonstrate that gene flow across ploidies appears to occur range-wide where *H. versicolor* and *H. chrysoscelis* are sympatric. Furthermore, the model support from our ABC analysis as well as the lack of the *H. versicolor* specific cluster in sympatric *H. chrysoscelis* populations provides strong evidence that this introgression is unidirectional with diploids contributing alleles to tetraploid populations but not *vice-versa*. Finally, our ILS assessment demonstrates a similar pattern to the parallel allele frequencies reported in Ralin et al. (1983) and Romano et al. (1987), but our results suggest the parallel pattern they observed were likely due to gene flow from *H. chrysoscelis* into *H. versicolor*, rather than parallel evolution.

As evidenced here, this reticulate history has led to three general lineages of *H. versicolor* that are distinguished by their nuclear and mitochondrial genomic compositions. Results from our mitochondrial divergence date estimation and the posterior distribution from our ABC analysis places the WGD and origin of *H. versicolor* from extinct NE *H. chrysoscelis* as occurring sometime within the last 426,000 years, with the best estimates from the mitochondrial phylogenetic and ABC analyses at 262,000 and 100,000 years ago respectively. These dates place the WGD event on either side of the Illinois glacial period (approximately 190–130 Ka, Curry et al. 2011) and agrees with the previous estimate of other Pre-Wisconsin (roughly 150,000) years ago from (Blair 1965; Ralin and Selander 1979; Maxson et al. 1977). Coalescent times for MW *H. versicolor* from our mitochondrial phylogenetic analysis largely overlap with the NE *H. versicolor* coalescent time, suggesting that backcrossing of NE *H. versicolor* with the extinct MW *H. chrysoscelis* to form the MW *H. versicolor* lineage occurred shortly after the initial WGD event. Regarding SW *H. versicolor*, there is not a monophyletic mitochondrial clade for SW *H. versicolor* individually that allows for a date of the initial backcrossing of that group. However, evidence from this study suggests this lineage is the youngest of three major *H. versicolor* lineages. It is unclear whether the lack of monophyly is a result of mitochondria consistently migrating from *H. chrysoscelis* to SW *H. versicolor* or that differences in mitochondrial genomes between *H. chrysoscelis* and *H. versicolor* here are so small as to preclude phylogenetic resolution—though there is no evidence of mitochondrial introgression in MW and NE *H. versicolor* despite strong evidence of hybridization between *H. chrysoscelis* and *H. versicolor* in these populations. We do however see one other case of recent mitochondrial introgression in *H. versicolor,* wherein a single, highly disjunct population of *H. versicolor* in Meade county Kentucky has integrated novel mitochondria from Central *H. chrysoscelis* (CC) into at least one individual (Ind. 20; Supp. Table 1). While it is possible this disjunct population of *H. versicolor* is from a unique WGD event, the presence of *H. versicolor* specific clusters from STRUCTURE as well as our RAxML concatenated phylogenetic analyses instead suggest this population became separated from already established *H. versicolor* populations (Fig. 5b; Fig. 4c,d,f; Supp. Fig. 7).

Results from this study suggest there are three lineages of *H. chrysoscelis* that each encompass large geographic areas across eastern North America. While previous studies have recognized Eastern and Western *H. chrysoscelis* lineages with populations on the periphery of the two having an intermediate genetic makeup, our study instead suggests that intermediate *H. chrysoscelis* are a separate, paraphyletic Central *H. chrysoscelis* lineage that encompasses a monophyletic Eastern *H. chrysoscelis* lineage (Fig. 4b; Fig. 5c), and that the combined monophyletic clade of Eastern and Central *H. chrysoscelis* is sister to Western *H. chrysoscelis*. There is also evidence for ongoing hybridization between the lineages that overlap, particularly between Eastern and Central *H. chrysoscelis* in the Florida panhandle and between Central and Western *H. chrysoscelis* in southern Missouri. The case of hybridization in southern Missouri is particularly interesting, as our results suggest two diploid and two tetraploid lineages are all in contact and hybridizing in this area.

### Hyla versicolor in Comparison to Other Polyploid Systems

Though there are significant barriers to introgression across species of different ploidies, the high levels of hybridization observed in this study are not unusual. Among plants, hybridization between species and populations of different ploidies appears to occur regularly (Soltis and Soltis 2009; Alix et al. 2017). In anurans, the complexes of *Phyllomedusa tetraploidea*, *Odontophrynus americanus*, and *Bufo viridis* show strong evidence that hybridization between tetraploids and diploids regularly produces triploid individuals (Stöck et al. 2002, 2005, 2010; Brunes et al. 2010, 2015; Grenat et al. 2018), and there is some evidence that several populations of the latter complex are entirely composed of hybrid triploids that are now sexually reproducing (Stöck et al. 2002, 2012). Similarly, the Australian *Neobatrachus* complex is composed of several autopolyploid tetraploid and diploid species who have each likely hybridized with multiple species several times since the initial polyploids were formed (Novikova et al. 2020; Mable and Roberts 1997). In *Xenopus*, though there are no extant diploids with which polyploids can hybridize, polyploid species have regularly hybridized in the past, and this hybridization has resulted in the evolution of species with elevated ploidy levels ranging from 4n to 12n (Evans 2008). Outside of bisexual polyploid amphibians, in more unique systems such as the unisexual *Ambystoma laterale* system and the hybridogenetic *Pelophylax/Rana esculentus* complexes, hybridization is a *de facto* requirement for their existence. As suggested by Bogart and Bi (2013), hybridization across polyploids and with their lower-ploidy progenitors seems to be a necessary requirement for a polyploid species’ persistence. As outlined here, the *H. versicolor* complex does not appear to be an exemption from this claim but rather adds considerable support for its validity.

Finally, an important result from this study was the support for a singular origin of *H. versicolor.* Multiple origins are considered common in polyploidy, with some suggesting this is the rule rather than the exception (Soltis et al. 2003). Although this claim may be supported given the current synthesis of polyploid research, results from this study suggest studies concluding multiple origins have to be re-evaluated (see also, Arnold et al. 2015). Using this system as an example, the presence of several paraphyletic mitochondrial lineages associated with unique allelic variants was previously used as evidence that *H. versicolor* originated independently multiple times (Ptacek et al. 1994; Holloway et al. 2006), and similar evidence appears to be consistently used to support multiple origins in other systems (Segraves et al. 1999; Soltis et al. 2003, 2014). Our results suggest that simply the observation of patterns in the data that appear to support multiple origins of a polyploid species is in itself, insufficient evidence for multiple origins, and that for systems where there are patterns that support multiple origins, explicitly testing this hypothesis is necessary before drawing any definite conclusions.

### Conclusions

In the present study, we have outlined the evolutionary history of *H. versicolor* and *H. chrysoscelis* in relation to polyploid formation as suggested by the data and analyses at hand. Although any single method of analysis in polyploid systems is subject to limitations, our work demonstrates multiple lines of evidence, along with direct model testing of polyploid histories, is necessary to uncover the evolutionary origins of a polyploid complex. Our study has addressed several key questions but also identified several puzzling new patterns in this complex that merit additional research. Although extant examples of polyploids are rare in animals compared to other groups such as angiosperms, polyploidy and gene/chromosomal duplications have undoubtedly played a major role in the evolutionary history of many animal groups (Otto and Whitton 2000; Gregory and Mable 2005; Blomme et al. 2006). Polyploid evolution in animals, however, has received little attention compared to the body of work in plants. Here, we demonstrate with the gray treefrog complex that animal polyploids can provide intriguing systems with which to answer important questions regarding the origins of polyploidization and the consequences of this process for diversification and speciation.

## Acknowledgements

I would like to thank Anna Liner Farmer, Alisha Holloway, Mitch Tucker, David Cannatella, Travis LaDuc, and the Texas Biodiversity Center at the University of Texas, Austin for tissue and loans; Michelle Kortyna, Sean Holland, and Kirby Birch at FSU’s Center for Anchored Phylogenomics and Roger Mercer at FSU’s College of Medicine for laboratory support; Hank Bass, Jim Bogart, David Hillis, and Joseph Travis for helpful feedback during project development; Camille Roux, and Thibault Leroy for assistance with the ABC analysis. This work was supported by the National Science Foundation through an NSF GRF (DGE 1449440) and NSF DEB (1120516), and by the University of Missouri Research Council (URC-15-106).

## SUPPLEMENTARY INFORMATION

### Supplementary Methods

#### Anchored Hybrid Enrichment Bioinformatics Extended

In order to increase read accuracy and length, paired reads were merged prior to assembly following Rokyta et al. (2012). In short, for each degree of overlap each read pair was evaluated with respect to the probability of obtaining the observed number of matches by chance. The overlap with the lowest probability was chosen if the p-value was less than 10^-10^. This low p-value avoids chance matches in repetitive regions. Read pairs with a p-value below the threshold were merged and quality scores were recomputed for overlapping bases (Rokyta et al. 2012). Read pairs failing to merge were utilized but left unmerged during the assembly.

##### Read assembly

Divergent reference assembly was used to map reads to the probe regions and extend the assembly into the flanking regions (Prum et al. 2015). More specifically, a subset of the taxa used during probe design were chosen as references for the assembly: *Pseudacris nigrita* (Hylidae) and *Gastrophryne carolinensis* (Microhylidae). Matches were called if 17 bases matched a library of spaced 20-mers derived from the conserved reference regions (i.e., those used for probe design). Preliminary reads were then considered mapped if the 55 matches were found over 100 consecutive bases in the reference sequences (all possible gap-free alignments between the read and the reference were considered). The approximate alignment position of mapped reads were estimated using the position of the spaced 20-mer, and all 60-mers existing in the read were stored in a hash table used by the de-novo assembler. The de-novo assembler identifies exact matches between a read and one of the 60-mers found in the hash table. Simultaneously using the two levels of assembly described above, the three read files were traversed repeatedly until a pass through the reads produced no additional mapped reads. For each locus, a list of all 60-mers found in the mapped reads was compiled and 60-mers were clustered if found together in at least two reads. Contigs were estimated from 60-mer clusters. In the absence of contamination, low coverage, or gene duplication each locus should produce one assembly cluster. Consensus bases were called from assembly clusters as unambiguous base calls if polymorphisms could be explained as sequencing error (assuming a binomial probability model with the probability of error equal to 0.1 and alpha equal to 0.05). Otherwise ambiguous bases were called following IUPAC codes (e.g., ‘R’ was used if ‘A’ and ‘G’ were observed). Called bases were soft-masked (made lowercase) for sites with coverage lower than 5. Assembled contigs derived from less than 400 reads were removed to reduce the effects of cross contamination and rare sequencing errors in index reads (Assembly code available at doi:10.5061/dryad.51v22).

##### Allele phasing

Using the Bayesian approach developed by Pyron et al. (2016), we took the identified polymorphic sites within each assembly and used the read overlap information to determine the posterior distribution of allele phasing under assumptions of diploidy and separately under the assumption of tetraploidy. Markov chains were run for 20,000 generations, sampling every 100 generations after the first 10,000 generates were discarded as burnin. Haplotypes for each individual locus combination were identified by drawing a sample from the posterior distribution for the relevant assembly.

##### Orthology assessment

After grouping homologous sequences obtained by enrichment and whole genomes, orthology was determined for each locus following (Hamilton et al. 2016; Prum et al. 2015). For each locus, pairwise distances among homologs were computed using an alignment-free approach based on percent overlap of continuous and spaced 20-mers. Using the distance matrix, sequences were clustered using a Neighbor-Joining algorithm, but allowing at most one sequence per species to be in a given cluster. Note that flanks recovered through extension assembly contain more variable regions and allow gene copies to be sorted efficiently. Gene duplication before the basal ancestor of the clade results in two distinct clusters that are easily separated. Duplication within the clade typically results in two clusters, one containing all of the taxa and a second containing a subset of the taxa (missing data). Gene loss also results in missing data. In order to reduce the effects of missing data, clusters containing fewer than 50% of the species were removed from downstream processing.

##### Alignment and trimming

Orthologous sequences were processed using a combination of automated and manual steps in order to generate high quality alignments in a reasonable amount of time. Sequences in each orthologous cluster were aligned using MAFFT v7.023b (Katoh et al. 2002; Katoh and Standley 2013), with --genafpair and --maxiterate 1000 flags utilized. The alignment for each locus was then trimmed/masked using the steps from Prum et al. (2015). First, each alignment site was identified as “conserved” if the most commonly observed character was present in *>*50% of the sequences. Second, using a 20bp window, each sequence was scanned for regions that did not contain at least 12 characters matching to the common base at the corresponding conserved site. Characters from regions not meeting this requirement were masked. Third, sites with fewer than 32 unmasked bases were removed from the alignment. A visual inspection of each masked alignment was carried out in Geneious version 7. Regions of sequences identified as obviously misaligned or paralogous were removed. SNPs were extracted from alignments using a custom Java script that identified polymorphic sites flanked on either side by 5 base pairs that were conserved across all taxa.

#### Polyploid Speciation Model Testing Parameters

The framework for simulating data for our ABC analysis allows for 45 total possible models given the same prior distributions of parameters. The 45 models are possible given three modes of polyploidization, three chromosomal inheritance patterns, and 5 different migration histories. The three polyploidization modes and their parameters that could be simulated under this framework are: Allopolyploidization (two ancestral lineages split at time Tsplit, combined to form a polyploid at time Twgd), Autopolyploidization (Two lineages split at time Tsplit, one lineage splits into two subgenomes at time Twgd; in effect, this is autopolyploidization of a sister lineage to the sampled diploid lineage), and Autopolyploid Speciation (lineages and subgenomes both split from a single ancestral population at time Tsplit = Twgd; in effect, this is autpolyploidization of the sampled diploid lineage). Chromosomal inheritance is simulated by allowing either no migration across subgenomes (disomic inheritance), 10 migrants per generation between subgenomes across all loci (tetrasomic inheritance), or 10 migrants per generation between subgenomes for a random number of loci (heterosomic inheritance). Ten migrants were chosen as this is sufficient to prevent differentiation between the subgenomes at the chosen locus (Templeton 2006; Roux and Pannell 2015). Finally, migration was simulated as occurring either asymmetrically between both diploids and tetraploids or in a single direction with only the tetraploid receiving migrants from the diploid. Migration patterns can also be divided subgenomically, with migration occurring either between the diploid and both subgenomes (migAB, Fig. 2) of the tetraploid or between the diploid and only one subgenome (migA, Fig. 2). The number of migrants in any one simulation was generated by sampling from a beta distribution, who’s parameters themselves were drawn from a uniform distribution. Prior distributions for these parameters were: scalar (0,10), *α* (0,10), and *β* (0,100).

#### Use of the D3 Statistic for Polyploid Inference

In addition to summary statistics already reported in the literature, we also used a modified version of the D3 statistic (Hahn and Hibbins 2019) for our ABC analysis. The D3 statistic was developed as a metric to detect introgression using only three samples, comparing pairwise distances of two sister lineages to a relative of those two lineages. In brief, significant deviations from a D3 value of zero likely indicates introgression between one of the ingroup lineages and the sister of the two ingroup lineages.

For use in our ABC analysis, we calculated the D3 statistic with the two tetraploid (*H. versicolor*) alleles as our ingroup lineages and a random diploid (*H. chrysoscelis*) allele as the outgroup lineage. Calculating the D3 statistic this way would allow us to distinguish if one *H. versicolor* allele was more similar to *H. chrysoscelis* —potentially indicating an allopolyploid hybird origin of *H. versicolor*. Additionally, differing chromosomal inheritance patterns should also affect the D3 statsitic—as increasing tetrasomic inheritance should erode any differences between the two tetraploid alleles and push D3 to zero. Finally, we calculated only the absolute value of every D3 value at each locus because we did not want to bias our results by having separated the MIN/MAX subgenomes (below referred to as D3a; ‘a’ referring to absolute value).

In addition to calculating this statistic to use in our ABC analysis, we were also interested in the behavior of this statistic under various heteroploidy migration histories. To evaluate the utility of this statistic, we conducted 1000 20-locus simulations using the same scripts, software, input files, and prior values as we did for the ABC model testing analysis. We reported the mean and standard deviation D3a value calculated for each 20 locus simulation, resulting in a final dataset of 1000 values for each of the models.

### Supplementary Results

#### D3a Statistic

Simulations of the D3a statistic under each of the possible models show that this statistic can be useful in several ways, particularly in conjunction with other statistics to distinguish among polyploid speciation, inheritance, and migration models. Overall, when there is no migration between diploids and tetraploids, the average and standard deviation D3a values are both near zero except under allopolyploid disomic and heterosomic models (Fig. 12,13). Migration between diploids and tetraploids tends to produce higher average D3a values—with bidirectional migration models having the highest D3a values. Similarly, the standard deviation of D3a values across loci also distinguishes between migration models in the same fashion. Increasing levels of disomic inheritance tends to elevate average D3a values across all models as well as the standard deviation of D3a for models of no migration or bidirectional migration. While there is a large amount of variation within specific models, our results suggest that the D3a statistic can be generally useful for informing migration models, and if migration models are known, can be informative for determining chromosomal inheritance patterns and distinguishing allo vs. autopolyploidy in some cases.

**Supplemental Table 1.** Locality and specimen details for samples used in this study. Bold specimens indicate samples used in our migrate-n analysis.

| Species | Map ID | Collab ID | State | County | Latitude | Longitude |
| --- | --- | --- | --- | --- | --- | --- |
| *Hyla andersonii* | NA | WED54451 | NJ | Burlington | 39.8559 | -74.6869 |
| *Hyla arenicolor* | NA | DCC3043 | AZ | Coconino | 35.0596 | -111.719 |
| *Hyla arenicolor* | NA | TJL 1286 | TX | Presidio | 29.9895 | -104.104 |
| *Hyla avivoca* | NA | MP710 | AL | Macon | 32.4253 | -85.7476 |
| *Hyla avivoca* | NA | DCC3857 | GA | Chatham | 31.9994 | -81.1196 |
| *Hyla avivoca* | NA | HCG21 | LA | Grant | 31.6944 | -92.5396 |
| *Hyla avivoca* | NA | H146 | LA | Jefferson | 29.8237 | -90.1402 |
| *Hyla avivoca* | NA | HCG84 | LA | Madison | 32.4437 | -91.2876 |
| *Hyla avivoca* | NA | MP607 | MS | Hinds | 32.2889 | -90.8041 |
| *Hyla avivoca* | NA | HCG28 | TN | Obion | 36.3469 | -89.1706 |
| *Hyla chrysoscelis* | 36 | MP816 | VA | Mecklenberg | 36.5929 | -78.3428 |
| *Hyla chrysoscelis* | 37 | MP 584 | VA | Charles City | 37.36 | -77.06 |
| ***Hyla chrysoscelis*** | **38** | **VA11** | **VA** | **Halifax** | **36.717** | **-78.8042** |
| ***Hyla chrysoscelis*** | **39** | **MP002** | **GA** | **Chatham** | **32.0288** | **-81.2694** |
| *Hyla chrysoscelis* | 40 | MP 674 | VA | Charles City | 37.36 | -77.06 |
| ***Hyla chrysoscelis*** | **41** | **MY01** | **MD** | **Harford** | **39.512** | **-76.4358** |
| *Hyla chrysoscelis* | 42 | ECM7850 | FL | Leon | 30.4914 | -84.1134 |
| *Hyla chrysoscelis* | 43 | MP205 | SC | Jasper | 32.47 | -81.02 |
| *Hyla chrysoscelis* | 44 | MP273 | VA | Goochland | 37.67 | -77.84 |
| *Hyla chrysoscelis* | 45 | NC11 | NC | Orange | 35.9536 | -79.063 |
| ***Hyla chrysoscelis*** | **46** | **FL08** | **FL** | **Alachua** | **29.6556** | **-82.3272** |
| *Hyla chrysoscelis* | 47 | MP649 | GA | Houston | 32.24 | -83.37 |
| *Hyla chrysoscelis* | 48 | MP732 | FL | Leon | 30.479 | -84.235 |
| *Hyla chrysoscelis* | 49 | FL06 | FL | Alachua | 29.6728 | -82.3469 |
| *Hyla chrysoscelis* | 50 | FL72 | FL | Leon | 30.4379 | -84.2457 |
| ***Hyla chrysoscelis*** | **51** | **ECM4690** | **FL** | **Liberty** | **30.2409** | **-85.0151** |

| Species | Map ID | Collab ID | State | County | Latitude | Longitude |
| --- | --- | --- | --- | --- | --- | --- |
| *Hyla chrysoscelis* | 52 | ECM4533 | FL | Leon | 30.3249 | -84.4583 |
| ***Hyla chrysoscelis*** | **53** | **MP458** | **AL** | **Russell** | **32.22** | **-85.15** |
| *Hyla chrysoscelis* | 54 | MP139 | LA | East Baton Rouge | 30.3438 | -91.0865 |
| *Hyla chrysoscelis* | 55 | MP138 | LA | East Baton Rouge | 30.3438 | -91.0865 |
| ***Hyla chrysoscelis*** | **56** | **MP647** | **TX** | **Harrison** | **32.53** | **-94.35** |
| *Hyla chrysoscelis* | 57 | MP753 | KY | Carlisle | 36.8128 | -89.0082 |
| *Hyla chrysoscelis* | 58 | MP859 | GA | Upson | 32.9559 | -84.4616 |
| *Hyla chrysoscelis* | 59 | 14-1 | KY | Bell | 36.7281 | -83.7409 |
| *Hyla chrysoscelis* | 60 | MP692 | TN | Monroe | 35.5233 | -84.3628 |
| *Hyla chrysoscelis* | 61 | MP670 | FL | Okaloosa | 30.7529 | -86.6341 |
| *Hyla chrysoscelis* | 62 | MP263 | LA | St. Martins | 30.2211 | -91.958 |
| *Hyla chrysoscelis* | 63 | MP847 | LA | Lincoln | 32.5345 | -92.6899 |
| *Hyla chrysoscelis* | 64 | MP195 | LA | Allen | 30.67 | -92.79 |
| *Hyla chrysoscelis* | 65 | MP696 | TN | Shelby | 35.29 | -89.908 |
| *Hyla chrysoscelis* | 66 | MP249 | MS | Hinds | 32.27 | -90.42 |
| ***Hyla chrysoscelis*** | **67** | **MP651** | **WV** | **Summers** | **37.5282** | **-80.9949** |
| *Hyla chrysoscelis* | 68 | MSC02 | MO | Cape Girardeau | 37.2403 | -89.4988 |
| *Hyla chrysoscelis* | 69 | ECM3053 | IL | Jackson | 37.7592 | -89.2706 |
| *Hyla chrysoscelis* | 70 | MP701 | VA | Smyth | 32.9476 | -84.4568 |
| *Hyla chrysoscelis* | 71 | MP 453 | KY | Meade | 37.93 | -86.05 |
| *Hyla chrysoscelis* | 72 | MP693 | LA | St. Martins | 30.2211 | -91.958 |
| *Hyla chrysoscelis* | 73 | DCC3751 | IN | Monroe | 39.187 | -86.5 |
| *Hyla chrysoscelis* | 74 | ECM4466 | KY | Graves | 36.8393 | -88.5264 |
| ***Hyla chrysoscelis*** | **75** | **DCC3883** | **AL** | **Bibb** | **32.9563** | **-87.1423** |
| *Hyla chrysoscelis* | 76 | INHS131T | IL | Edgar | 39.7734 | -87.6847 |
| *Hyla chrysoscelis* | 77 | DCC3829 | OH | Ross | 39.395 | -82.9432 |
| *Hyla chrysoscelis* | 78 | MP630 | KY | Laurel | 37.13 | -84.1 |

| Species | Map ID | Collab ID | State | County | Latitude | Longitude |
| --- | --- | --- | --- | --- | --- | --- |
| *Hyla chrysoscelis* | 79 | MP 454 | KY | Meade | 37.93 | -86.05 |
| *Hyla chrysoscelis* | 80 | MP452 | KY | Meade | 37.93 | -86.05 |
| *Hyla chrysoscelis* | 81 | ECM4330 | IL | Jefferson | 38.15 | -88.847 |
| *Hyla chrysoscelis* | 82 | MP724 | MS | Hancock | 30.2625 | -89.5424 |
| ***Hyla chrysoscelis*** | **83** | **MP350** | **MO** | **Phelps** | **37.7** | **-91.89** |
| *Hyla chrysoscelis* | 84 | MP386 | MO | Howell | 36.77 | -91.87 |
| *Hyla chrysoscelis* | 85 | P112 | MO | Phelps | 37.6184 | -91.9983 |
| *Hyla chrysoscelis* | 86 | 14-6 | MO | Phelps | 37.6187 | -92.0004 |
| *Hyla chrysoscelis* | 87 | MP370 | MO | Oregon | 36.69 | -91.41 |
| ***Hyla chrysoscelis*** | **88** | **14-7** | **MO** | **Phelps** | **37.6187** | **-92.0004** |
| *Hyla chrysoscelis* | 89 | DCC3874 | IA | Butler | 42.7524 | -92.8562 |
| ***Hyla chrysoscelis*** | **90** | **MP706** | **IA** | **Clarke** | **41.029** | **-93.4506** |
| ***Hyla chrysoscelis*** | **91** | **MP773** | **MO** | **Barry** | **36.8838** | **-93.6648** |
| *Hyla chrysoscelis* | 92 | MP717 | TX | Smith | 32.6334 | -95.3569 |
| *Hyla chrysoscelis* | 93 | MP804 | NE | Otoe | 40.7272 | -96.1212 |
| *Hyla chrysoscelis* | 94 | MP723 | TX | Bastrop | 30.0797 | -97.238 |
| *Hyla chrysoscelis* | 95 | T-1 | KS | Douglas | 38.9181 | -95.2311 |
| *Hyla chrysoscelis* | 96 | INHS924T | IL | Ogle | 42.0431 | -89.3841 |
| *Hyla chrysoscelis* | 97 | MP327 | OK | Payne | 36.1 | -96.99 |
| *Hyla chrysoscelis* | 98 | MP632 | MN | Hennepin | 45.0293 | -93.5148 |
| ***Hyla chrysoscelis*** | **99** | **MP802** | **MN** | **Ottertail** | **46.4265** | **-95.5577** |
| *Hyla chrysoscelis* | 100 | 14-5 | MI | Schoolcraft | 46.2895 | -85.9476 |
| *Hyla chrysoscelis* | 101 | MP135 | TX | Travis | 30.32 | -97.69 |
| *Hyla chrysoscelis* | 102 | MP329 | OK | Ottawa | 36.85 | -94.8 |
| *Hyla chrysoscelis* | 103 | MP686 | TX | Eastland | 32.3354 | -98.8294 |
| *Hyla chrysoscelis* | 104 | C10 | MN | Carver | 44.8789 | -93.7187 |
| ***Hyla chrysoscelis*** | **105** | **TX25** | **TX** | **Kendall** | **29.9794** | **-98.7231** |

| Species | Map ID | Collab ID | State | County | Latitude | Longitude |
| --- | --- | --- | --- | --- | --- | --- |
| *Hyla femoralis* |  | DCC3858 | GA | Chatham | 32.0054 | -81.122 |
| *Hyla versicolor* | 1 | MP702 | MN | Clearwater | 47.25 | -95.25 |
| ***Hyla versicolor*** | **2** | **MP795** | **MN** | **Ottertail** | **46.4265** | **-95.5577** |
| *Hyla versicolor* | 3 | DCC3864 | MN | St. Louis | 47.7681 | -92.3931 |
| ***Hyla versicolor*** | **4** | **MP099** | **OK** | **Cleveland** | **35.1595** | **-97.3985** |
| *Hyla versicolor* | 5 | MP045 | MO | Greene | 37.1868 | -93.1264 |
| ***Hyla versicolor*** | **6** | **INHS399T** | **IL** | **Iroquois** | **40.8791** | **-87.6876** |
| *Hyla versicolor* | 7 | MP759 | MO | Barry | 36.8838 | -93.6648 |
| *Hyla versicolor* | 8 | MP793 | OK | Ottawa | 36.8012 | -94.71 |
| *Hyla versicolor* | 9 | MP625 | Ontario | Guelph | 43.5497 | -80.1784 |
| *Hyla versicolor* | 10 | INHS950T | IL | Hancock | 40.5864 | -91.3065 |
| ***Hyla versicolor*** | **11** | **DCC3832** | **PA** | **Crawford** | **41.5667** | **-80.45** |
| ***Hyla versicolor*** | **12** | **DCC3828** | **OH** | **Ross** | **39.395** | **-82.9432** |
| *Hyla versicolor* | 13 | DCC3768 | IN | Porter | 41.5167 | -87.0833 |
| ***Hyla versicolor*** | **14** | **MP019** | **TX** | **Bastrop** | **30.13** | **-97.41** |
| *Hyla versicolor* | 15 | MP359 | MO | Ozark | 36.7601 | -92.1537 |
| *Hyla versicolor* | 16 | MP524 | MO | Phelps | 37.7 | -91.89 |
| ***Hyla versicolor*** | **17** | **BCHV02** | **MO** | **Boone** | **38.7611** | **-92.1995** |
| ***Hyla versicolor*** | **18** | **MP409** | **MO** | **Howell** | **36.77** | **-91.87** |
| ***Hyla versicolor*** | **19** | **TXHV03** | **TX** | **Wood** | **32.5887** | **-95.3249** |
| *Hyla versicolor* | 20 | MP 825 | KY | Meade | 37.93 | -86.05 |
| ***Hyla versicolor*** | **21** | **MP700** | **TN** | **Shelby** | **35.29** | **-89.908** |
| *Hyla versicolor* | 22 | MP162 | LA | Allen | 30.67 | -92.79 |
| *Hyla versicolor* | 23 | MP678 | WV | Summers | 37.56 | -80.92 |
| ***Hyla versicolor*** | **24** | **DCC3787** | **NY** | **Rensselaer** | **42.55** | **-73.625** |
| *Hyla versicolor* | 25 | MP020 | VA | Giles | 37.3591 | -80.5353 |
| *Hyla versicolor* | 26 | DCC3807 | CT | Tolland | 41.8075 | -72.2621 |

| Species | Map ID | Collab ID | State | County | Latitude | Longitude |
| --- | --- | --- | --- | --- | --- | --- |
| ***Hyla versicolor*** | **27** | **MP576** | **ME** | **Penobscot** | **44.9024** | **-68.6598** |
| *Hyla versicolor* | 28 | DCC3795 | NY | Westchester | 41.2728 | -73.8942 |
| *Hyla versicolor* | 29 | DCC3800 | CT | Windham | 41.7903 | -71.9422 |
| ***Hyla versicolor*** | **30** | **DCC3817** | **MD** | **Prince Georges** | **39.0572** | **-76.816** |
| ***Hyla versicolor*** | **31** | **DCC3816** | **MD** | **Prince Georges** | **39.0572** | **-76.816** |
| ***Hyla versicolor*** | **32** | **MP296** | **VA** | **Goochland** | **37.67** | **-77.84** |
| *Hyla versicolor* | 33 | MYHV05 | MD | Harford | 39.512 | -76.4358 |
| *Hyla versicolor* | 34 | DCC3823 | MD | Ann Arundel | 39.0572 | -76.816 |
| *Hyla versicolor* | 35 | MP809 | VA | Mecklenberg | 36.5929 | -78.3428 |

**Supplemental Table 2.** Summary statistics used for the ABC analysis and the observed statistics estimated from 50 loci comparing Eastern *H. chrysoscelis* (A) Northeast*H. versicolor* (B). *p*-values were generated from randomization tests of the observed estimate for 50 loci against a null of 1000 50 random loci estimates.

| Statistic | Estimate (50 loci) | *p*-value |
| --- | --- | --- |
| bialsites_avg | 20.4600 | 0.694 |
| bialsites_std | 13.1411 | 0.564 |
| sf_avg | 0.00005 | 0.507 |
| sf_std | 0.00031 | 0.525 |
| sxA_avg | 0.00119 | 0.645 |
| sxA_std | 0.00125 | 0.590 |
| sxB_avg | 0.00986 | 0.432 |
| sxB_std | 0.00883 | 0.258 |
| ss_avg | 0.00309 | 0.693 |
| ss_std | 0.00262 | 0.627 |
| successive_ss_avg | 1.10000 | 0.499 |
| successive_ss_std | 1.44568 | 0.216 |
| piA_avg | 0.00152 | 0.606 |
| piA_std | 0.00124 | 0.302 |
| piB_avg | 0.00300 | 0.600 |
| piB_std | 0.00294 | 0.297 |
| pearson_r_pi | 0.55675 | 0.583 |
| thetaA_avg | 0.00158 | 0.737 |
| thetaA_std | 0.00117 | 0.562 |
| thetaB_avg | 0.00369 | 0.501 |
| thetaB_std | 0.00289 | 0.219 |
| pearson_r_theta | 0.52557 | 0.576 |
| DtajA_avg | -0.21521 | 0.178 |
| DtajA_std | 0.72769 | 0.738 |
| DtajB_avg | -0.83063 | 0.875 |
| DtajB_std | 0.71572 | 0.402 |
| divAB_avg | 0.00320 | 0.267 |
| divAB_std | 0.00436 | 0.116 |
| netdivAB_avg | 0.00094 | 0.138 |
| netdivAB_std | 0.00287 | 0.122 |
| minDivAB_avg | 0.00039 | 0.188 |
| minDivAB_std | 0.00227 | 0.228 |
| maxDivAB_avg | 0.00709 | 0.438 |
| maxDivAB_std | 0.00629 | 0.146 |
| Gmin_avg | 0.02343 | 0.316 |
| Gmin_std | 0.09375 | 0.429 |
| Gmax_avg | 2.93673 | 0.614 |
| Gmax_std | 1.37469 | 0.230 |
| FST_avg | 0.14819 | 0.458 |
| FST_std | 0.14471 | 0.061 |
| D3a_avg | 0.37378 | 0.851 |
| D3a_std | 0.11024 | 0.496 |
| pearson_r_divAB_netDivAB | 0.94684 | 0.079 |
| pearson_r_divAB_FST | 0.55624 | 0.333 |
| pearson_r_netDivAB_FST | 0.63705 | 0.572 |
| ss_sf | 0.02000 | 0.483 |
| ss_noSf | 0.86000 | 0.921 |
| noSs_sf | 0.02000 | 0.382 |
| noSs_noSf | 0.10000 | 0.244 |

**Supplemental Table 3.** Estimates of the average and standard deviation for Tajima’s *θ* (*π*), Watterson’s *θ*, and Tajima’s *D* across all loci in *H. chrysoscelis*, all *H. versicolor* with MIN/MAX sequences combined or separated, and in individual *H. versicolor* lineages with MIN/MAX sequences combined or separated.

|  | *H. chrysoscelis* | All_MIN_MAX | All_MIN | All_MAX | NE_MAX | NE_MIN | MW_MIN | MW_MIN | SW_MAX | SW_MIN |
| --- | --- | --- | --- | --- | --- | --- | --- | --- | --- | --- |
| pi_avg | 0.00214 | 0.00293 | 0.00193 | 0.00373 | 0.00359 | 0.00198 | 0.00357 | 0.00204 | 0.00353 | 0.00187 |
| pi_std | 0.00133 | 0.00261 | 0.00218 | 0.00285 | 0.00271 | 0.00223 | 0.00292 | 0.00225 | 0.00288 | 0.00235 |
| theta_avg | 0.00321 | 0.00476 | 0.0027 | 0.00491 | 0.00383 | 0.00222 | 0.00379 | 0.00219 | 0.00389 | 0.00209 |
| theta_std | 0.00160 | 0.00312 | 0.00256 | 0.00301 | 0.00266 | 0.00244 | 0.00295 | 0.00237 | 0.00292 | 0.00233 |
| Dtaj_avg | -1.08906 | -1.32586 | -0.97056 | -0.93785 | -0.33412 | -0.46295 | -0.32657 | -0.31600 | -0.52141 | -0.56075 |
| Dtaj_std | 0.60000 | 0.61249 | 0.69021 | 0.59352 | 0.60692 | 0.74537 | 0.68073 | 0.76209 | 0.62047 | 0.69027 |

**Supplemental Table 4.** Coalescent timing for clades of interest estimated from BEAST analysis. Colors correspond to clade and background colors in Fig. 5

| Clade | Fig. 5 Color | Mean | 95% CI |
| --- | --- | --- | --- |
| All *H. chrysoscelis*/*H. versicolor*/*H. avivoca* | NA | 1.80 Ma | 0.989-2.85 Ma |
| All *H. chrysoscelis* | NA | 1.26 Ma | 0.675-1.99 Ma |
| NE *H. versicolor*/East *H. avivoca* | Green | 1.51 Ma | 0.812-2.37 Ma |
| NE *H. versicolor* | Green | 0.262 Ma | 0.125-0.426 Ma |
| MW *H. versicolor* | Yellow | 0.338 Ma | 0.131-0.430 Ma |
| Western *H. chrysoscelis* | Orange | 0.262 Ma | 0.130-0.430 Ma |
| FL Eastern/Eastern/Central *H. chrysoscelis* | Purple | 0.811 Ma | 0.431-1.27 Ma |
| Eastern/Central *H. chrysoscelis* | Purple | 0.527 Ma | 0.271-0.826 Ma |
| West *H. avivoca*, SW *H. versicolor*, Central *H. chrysoscelis* (CSW) | Light Blue | 0.591 Ma | 0.326-0.939 Ma |
| SW *H. versicolor*, Central *H. chrysoscelis* (CSW) | Light Blue | 0.223 Ma | 0.012-0.360 Ma |

**Supplemental Figure 1.**
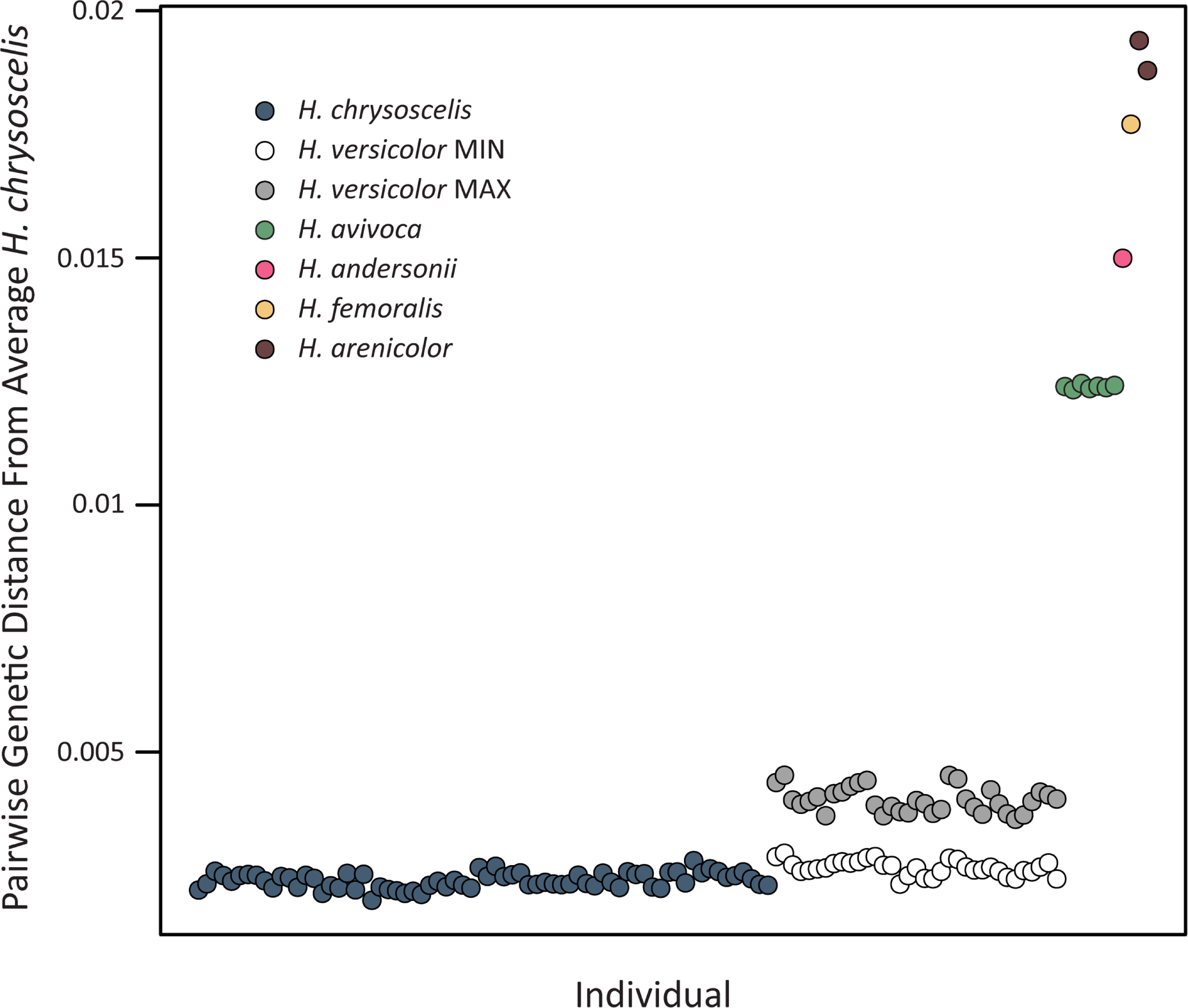
Pairwise genetic distance for each sampled individual to the average *H. chrysoscelis*. Averages were generated by calculating the pairwise genetic distance for each individual to each *H. chrysoscelis* (excluding themselves if the sample was *H. chrysoscelis*) and averaging over the total. *H. versicolor* individuals are separated into putative MIN/MAX subgenomes on the same x-axis plane.

**Supplemental Figure 2.**
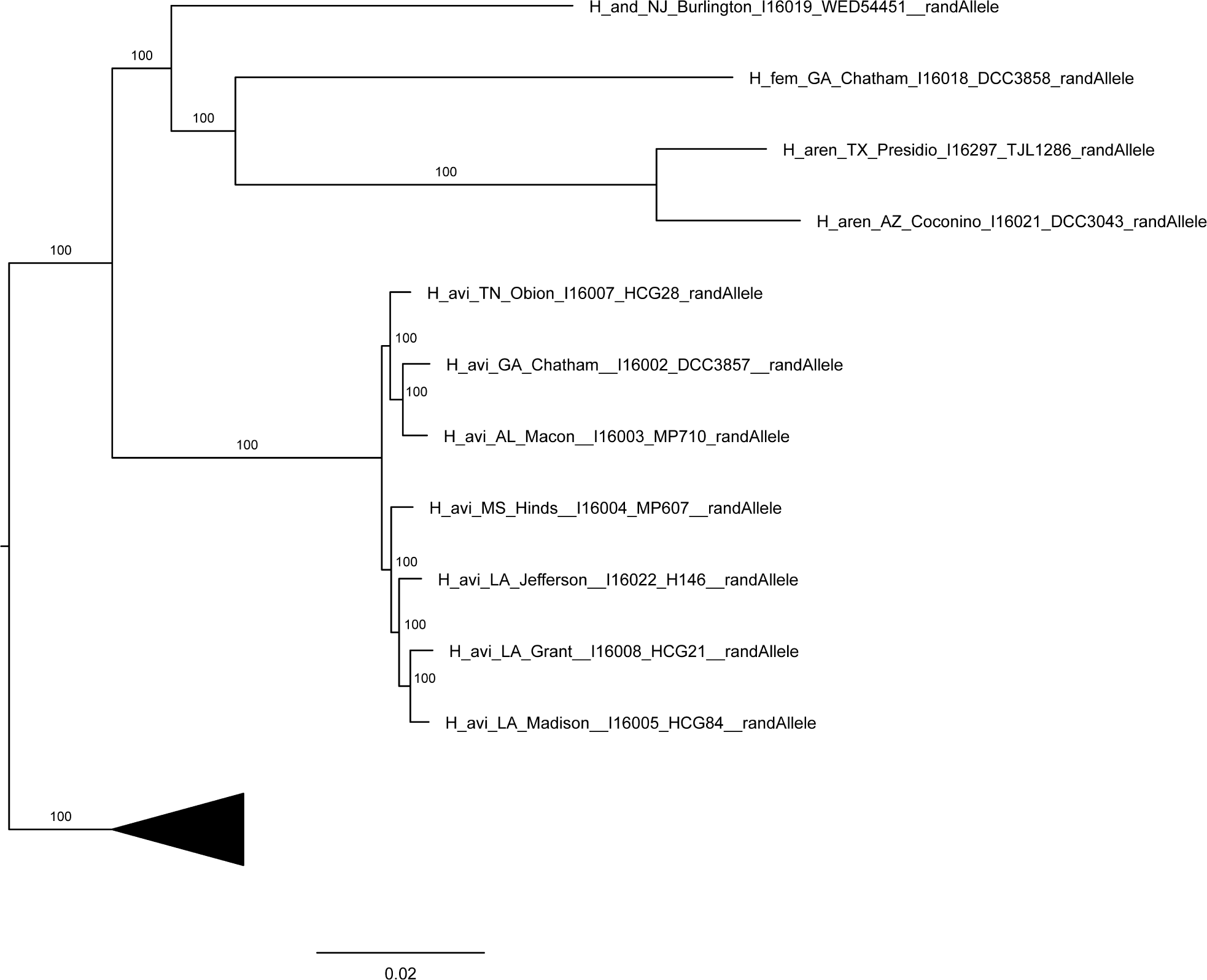
Nuclear phylogenetic relationships of outgroup taxa (*H. andersonii*, *H. arenicolor*, and *H. femoralis*) and *H. avivoca* relative to *H. versicolor* and *H. chrysoscelis* (collapsed) from the RAxML concatenated analysis using 244 AHE loci (Dataset 1 shown; all datasets produced the same relationships and support). Branch labels show bootstrap support values. Scale bar and branch lengths represent substitutions per site.

**Supplemental Figure 3.**
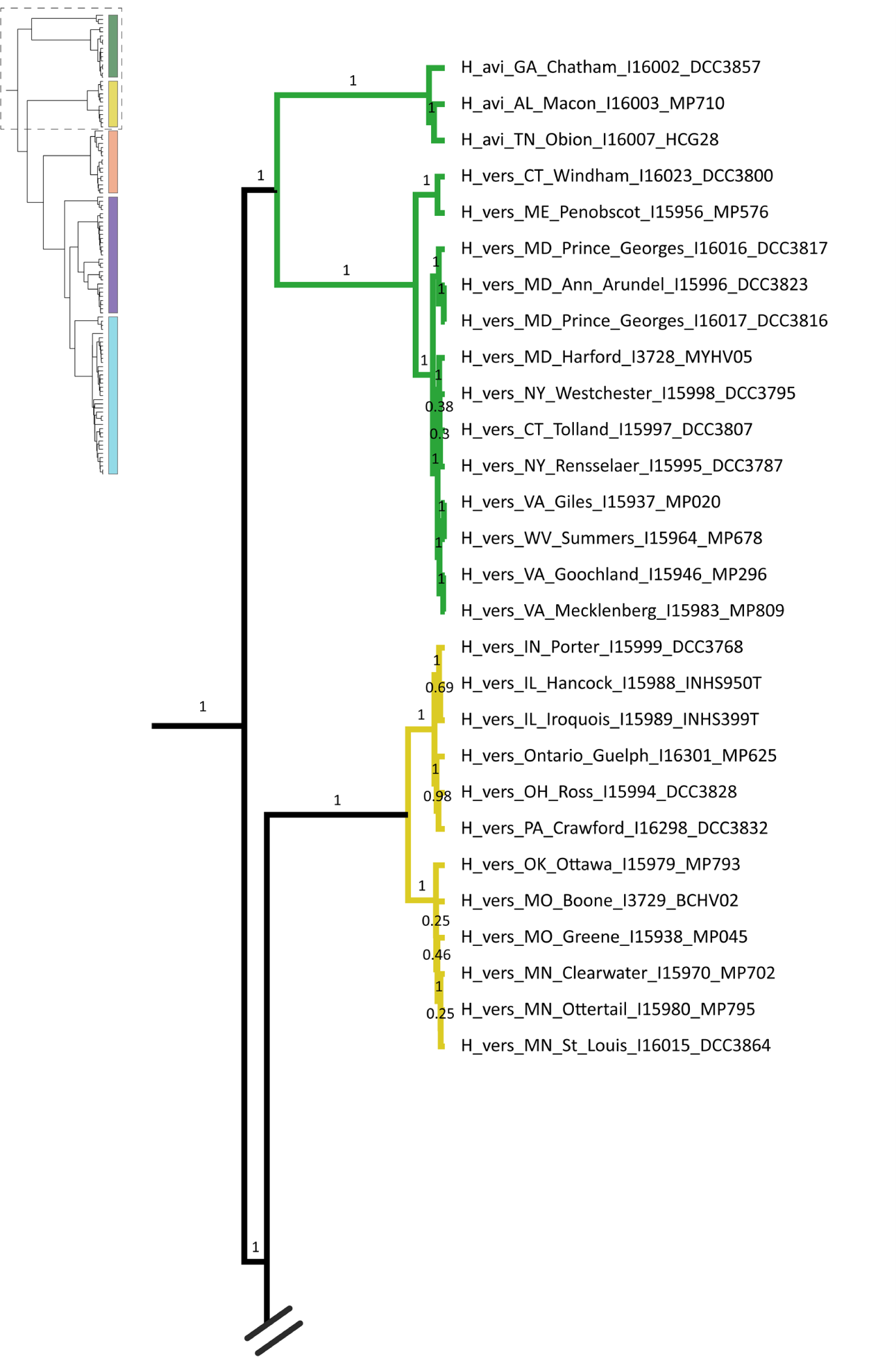
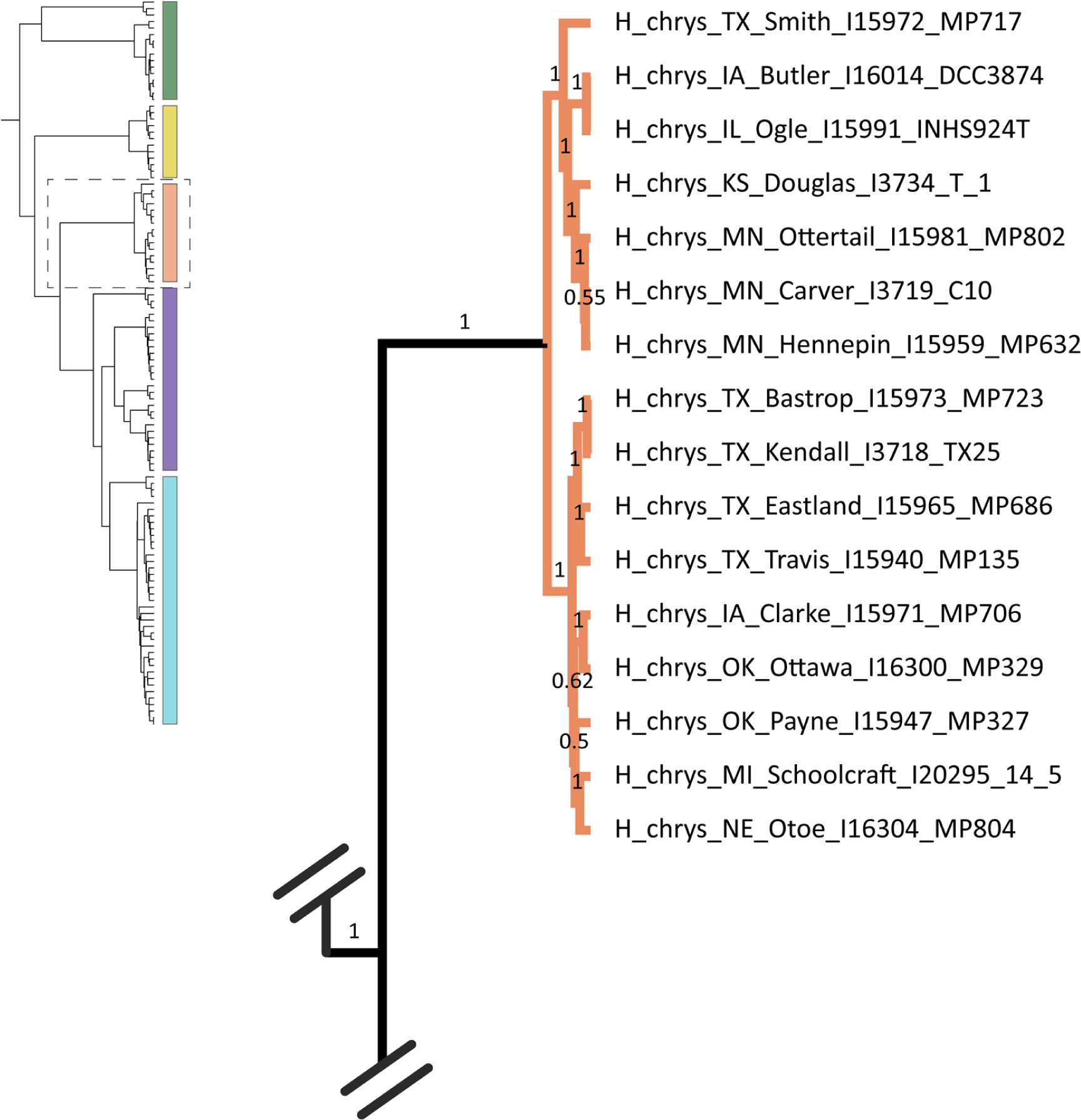
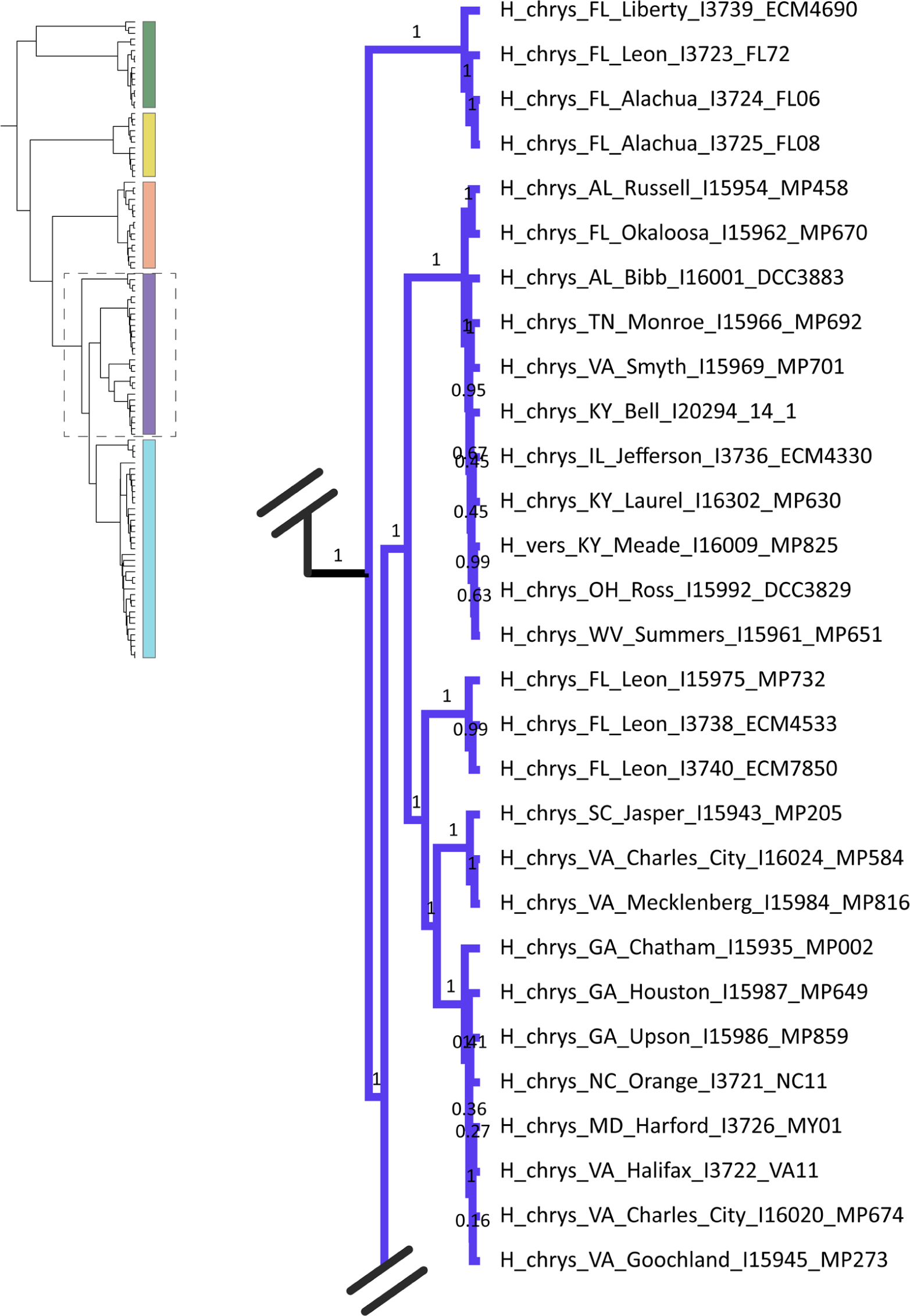
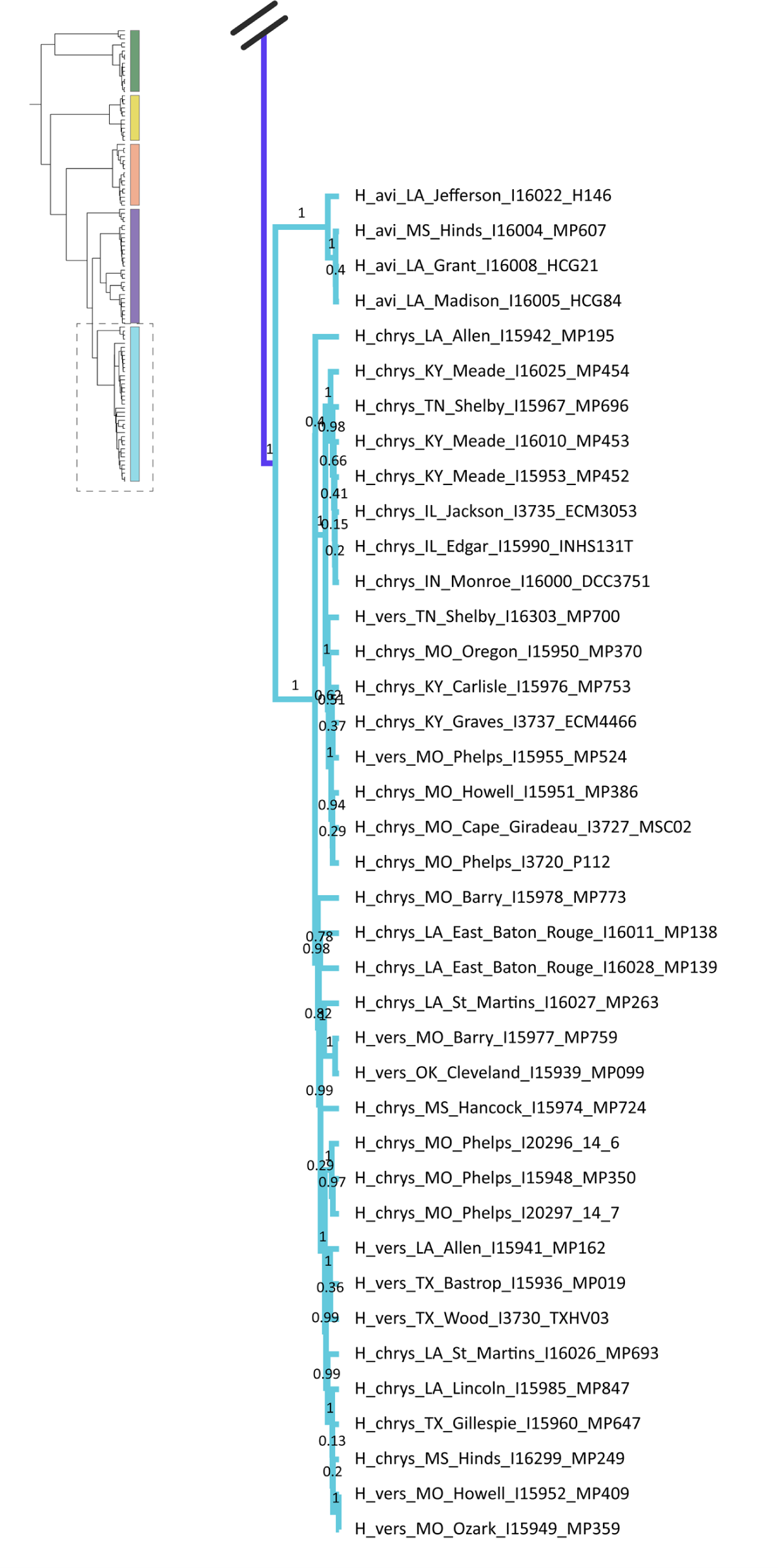
Expanded mitochondrial tree shown in Fig. 5a from the BEAST 2 analysis. Branch labels show posterior probability.

**Supplemental Figure 4.**
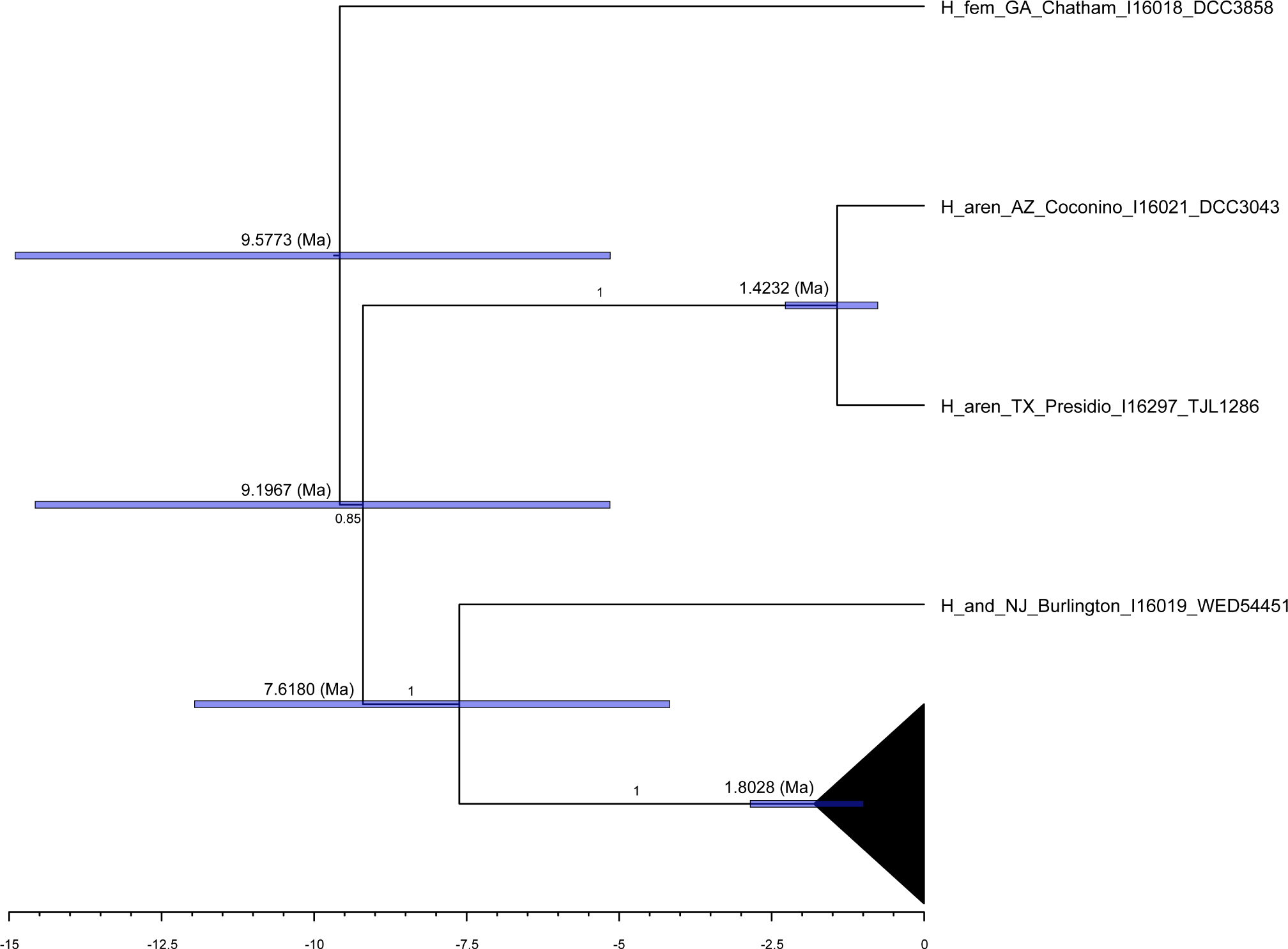
Mitochondrial relationships and coalescent timing of outgroup taxa (*H. andersonii*, *H. arenicolor*, and *H. femoralis*) relative to *H. avivoca*, *H. versicolor*, and *H. chrysoscelis* (collapsed) from the Beast 2 analysis. Branch labels show posterior probability. Node bars demonstrate the 95%CI estimated coalescent timing in millions of years (Ma), with the mean estimated coalescent time above the bars.

**Supplemental Figure 5.**
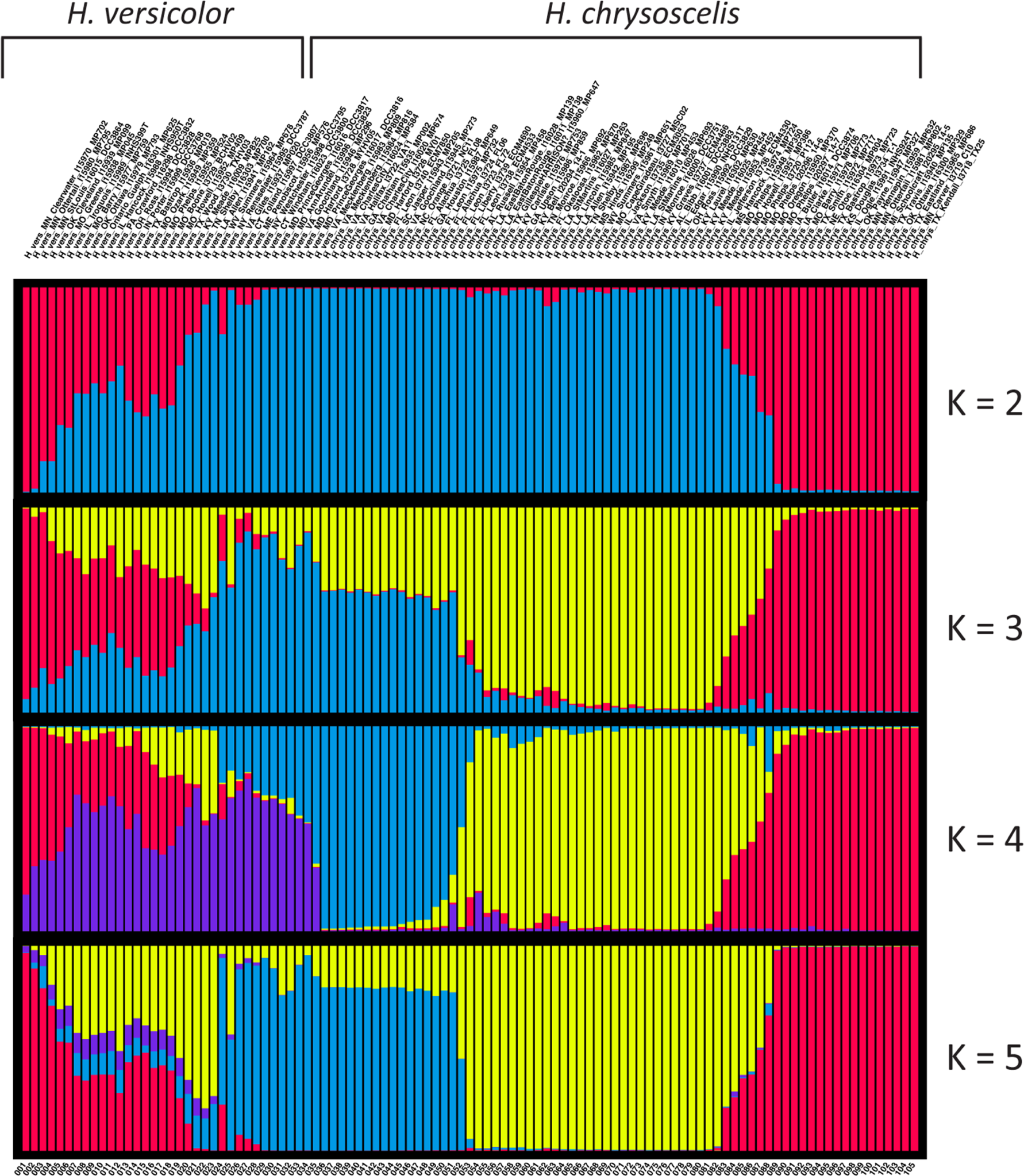
STRUCTURE results from an analysis of a single SNP per locus across 244 loci and including all *H. versicolor* and *H. chrysoscelis* samples. Pictured K values stop at 5, because clusters past k=4 were of a small proportion and did not appear to be concentrated across any biologically meaningful group.

**Supplemental Figure 6.**
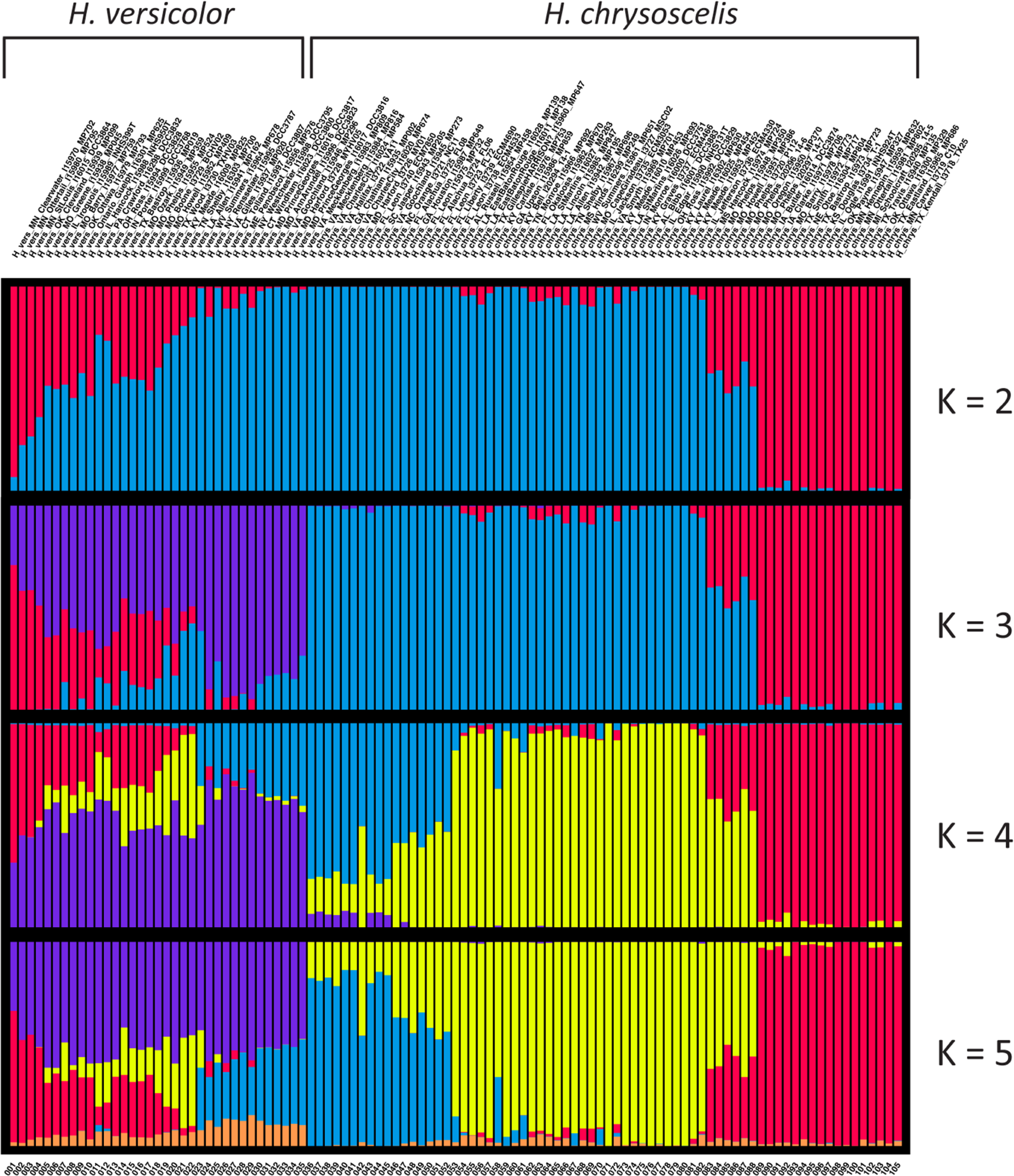
STRUCTURE results from an analysis of all 8,683 SNPs across 244 loci and including all *H. versicolor* and *H. chrysoscelis* samples. Pictured K values stop at 5, because clusters past k=4 were of a small proportion and did not appear to be concentrated across any biologically meaningful group.

**Supplemental Figure 7.**
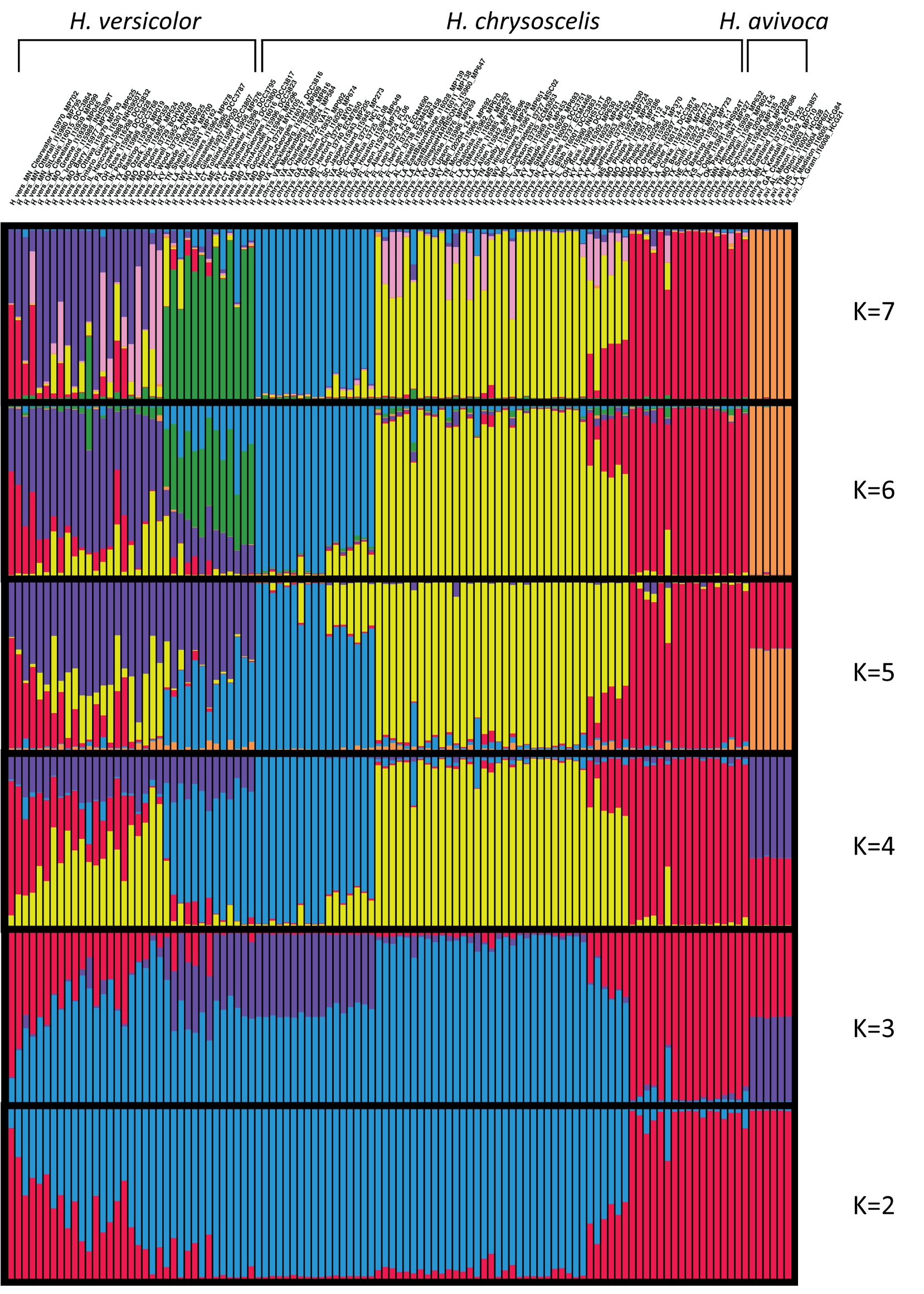
STRUCTURE results from an analysis of a single SNP per locus across 244 loci and including all *H. versicolor*, *H. chrysoscelis*, and *H. avivoca* samples. Order is the same as Fig. 5 and 6, with *H. avivoca* on the far right.

**Supplemental Figure 8.**
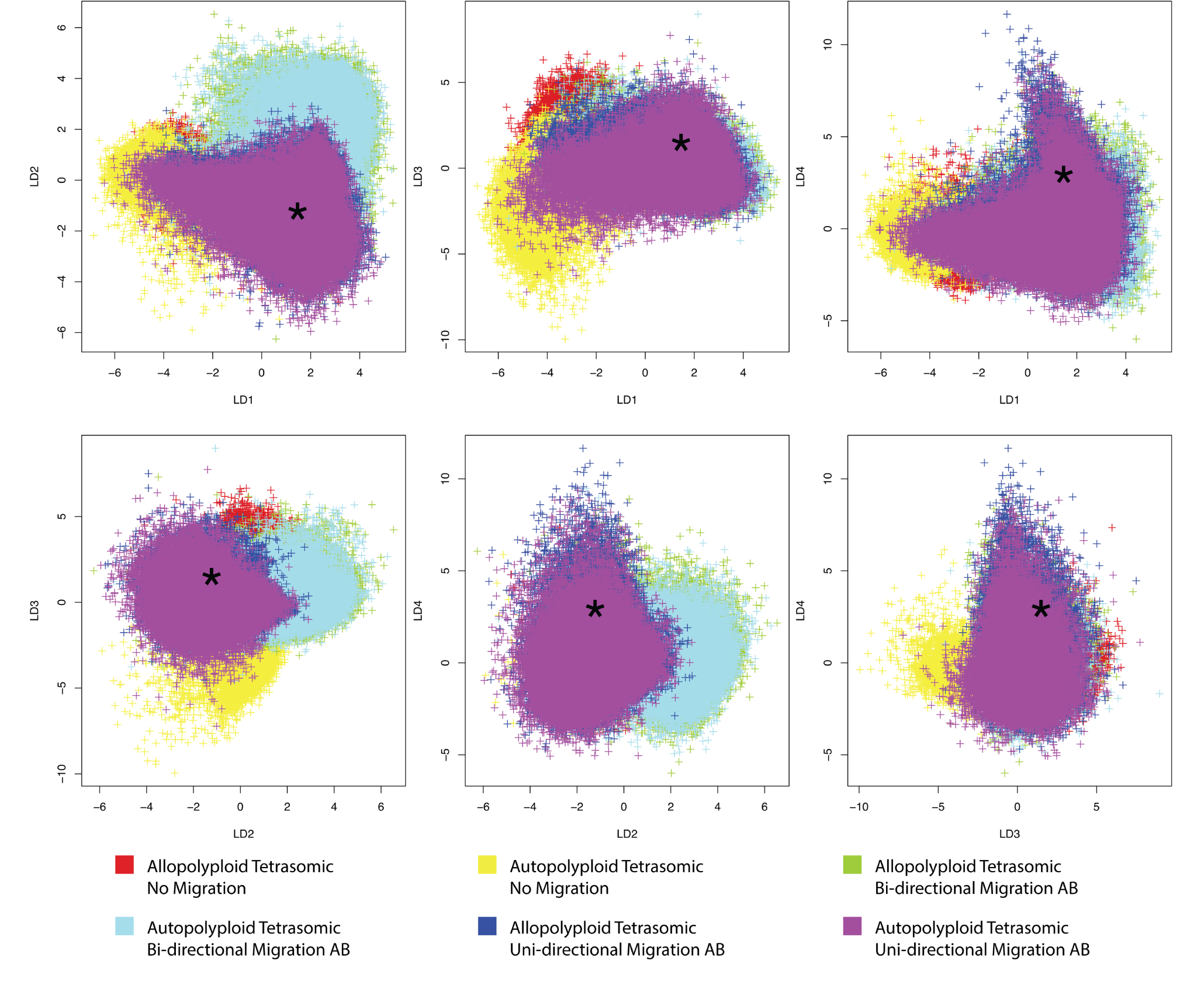
Pairwise comparison of LDA axes 1-4 for each of the 6 simulated polyploid speciation and migration models. The star represents the observed LDA values for the Northeast *H. versicolor* and Eastern *H. chrysoscelis* dataset.

**Supplemental Figure 9.**
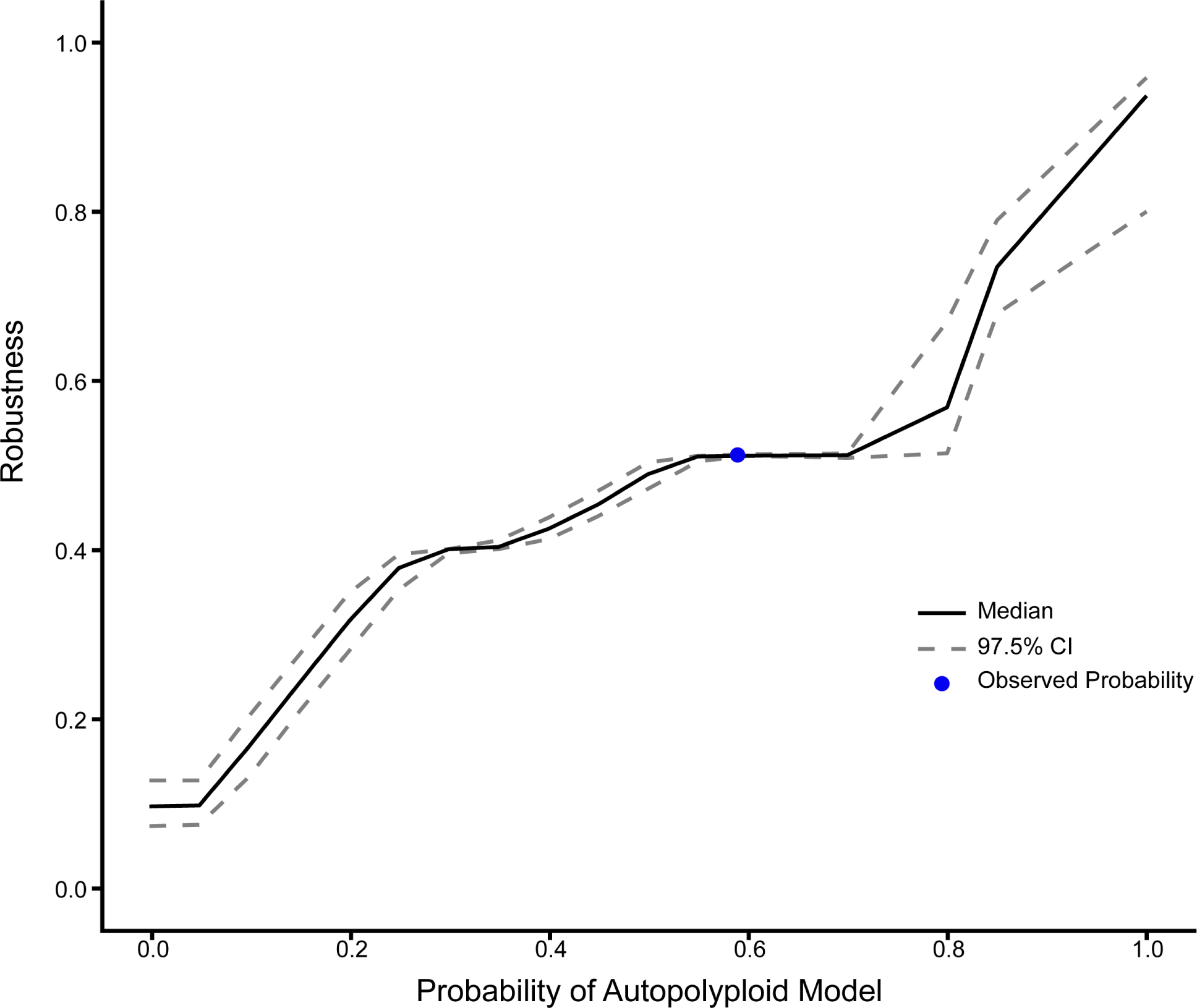
Robustness of our ABC analysis to select the true model (here, autopolyploid with tetrasomic inheritance and one-way AB migration) given an estimated probability of that model. Robustness was assessed using 1000 pseudo-observed datasets and calculated as 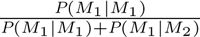. The blue circle shows the probability of the Autopolyploid model from our analysis of the observed data.

**Supplemental Figure 10.**
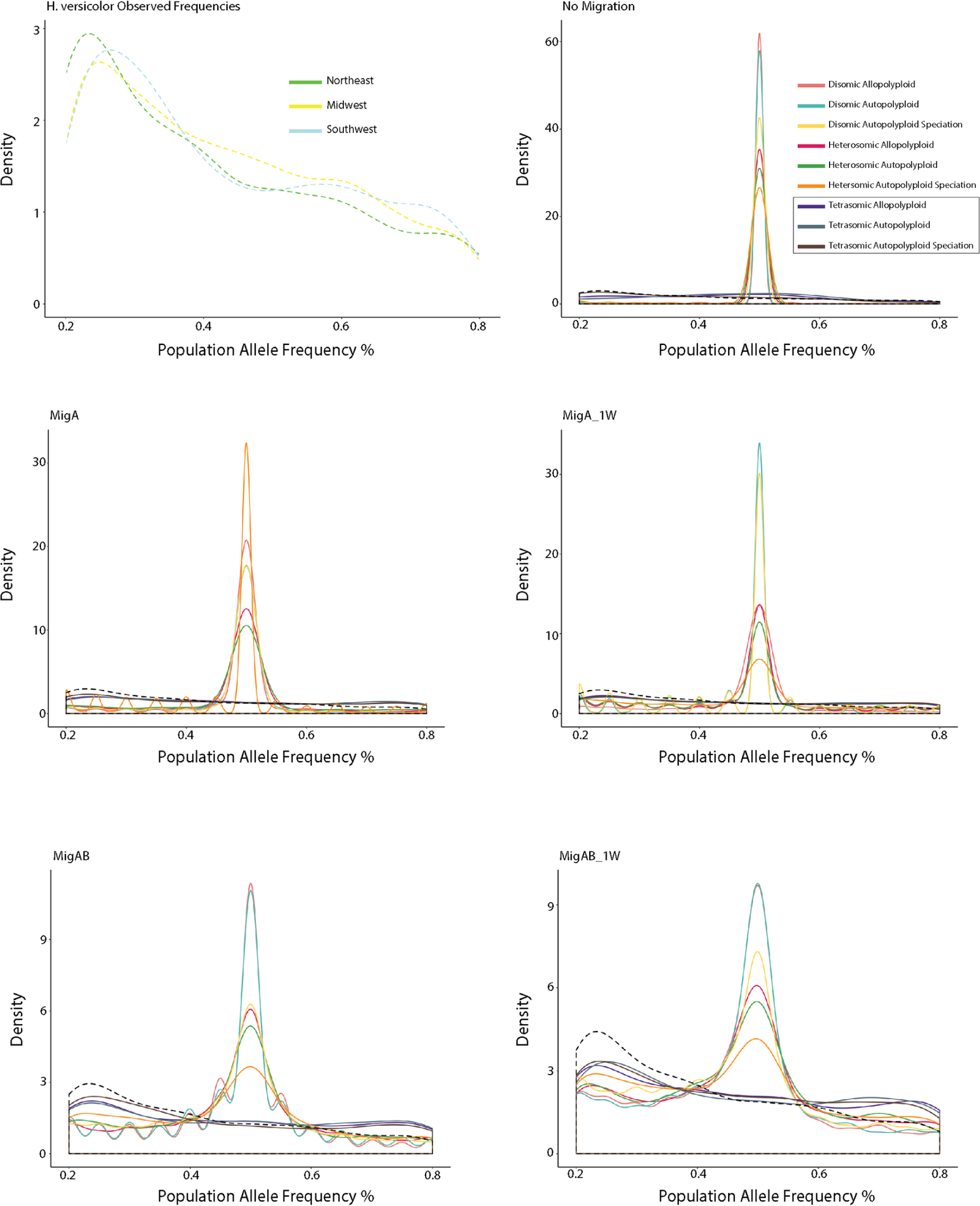
Observed and simulated population allele frequencies. First panel: Observed intermediate population allele frequency density plots for each *H. versicolor* mitochondrial lineage. Following panels: Observed and simulated population allele frequency density plots under each polyploid speciation and inheritance model separated by migration model. Simulated allele frequencies are solid lines, observed allele frequency of Northeast *H. versicolor* shown by the dashed line.

**Supplemental Figure 11.**
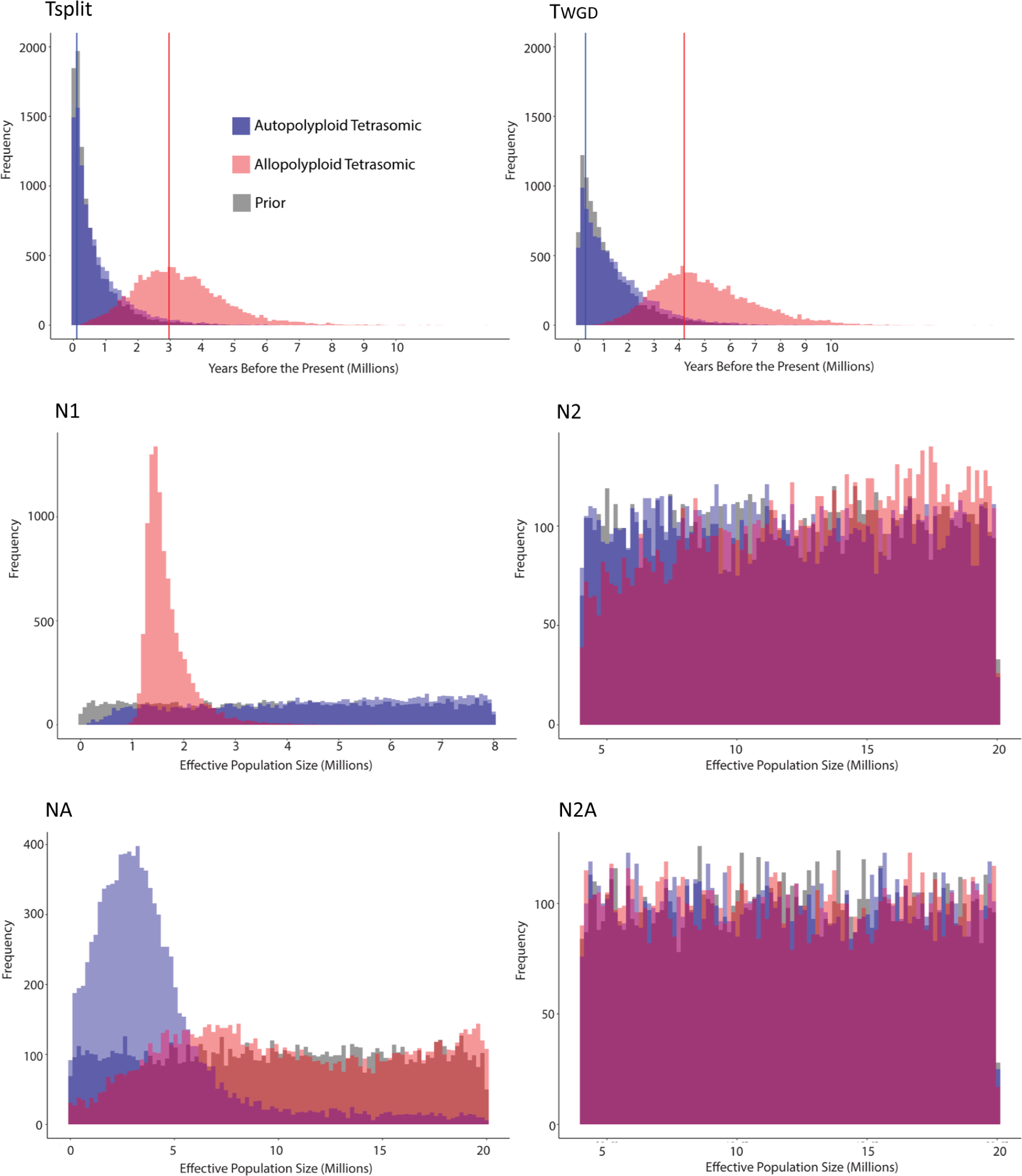
Prior and posterior distributions for the two best supported models from the ABC analysis. Both models presented are with a unidirectional migration history. Prior distribution is in gray.

**Supplemental Figure 12.**
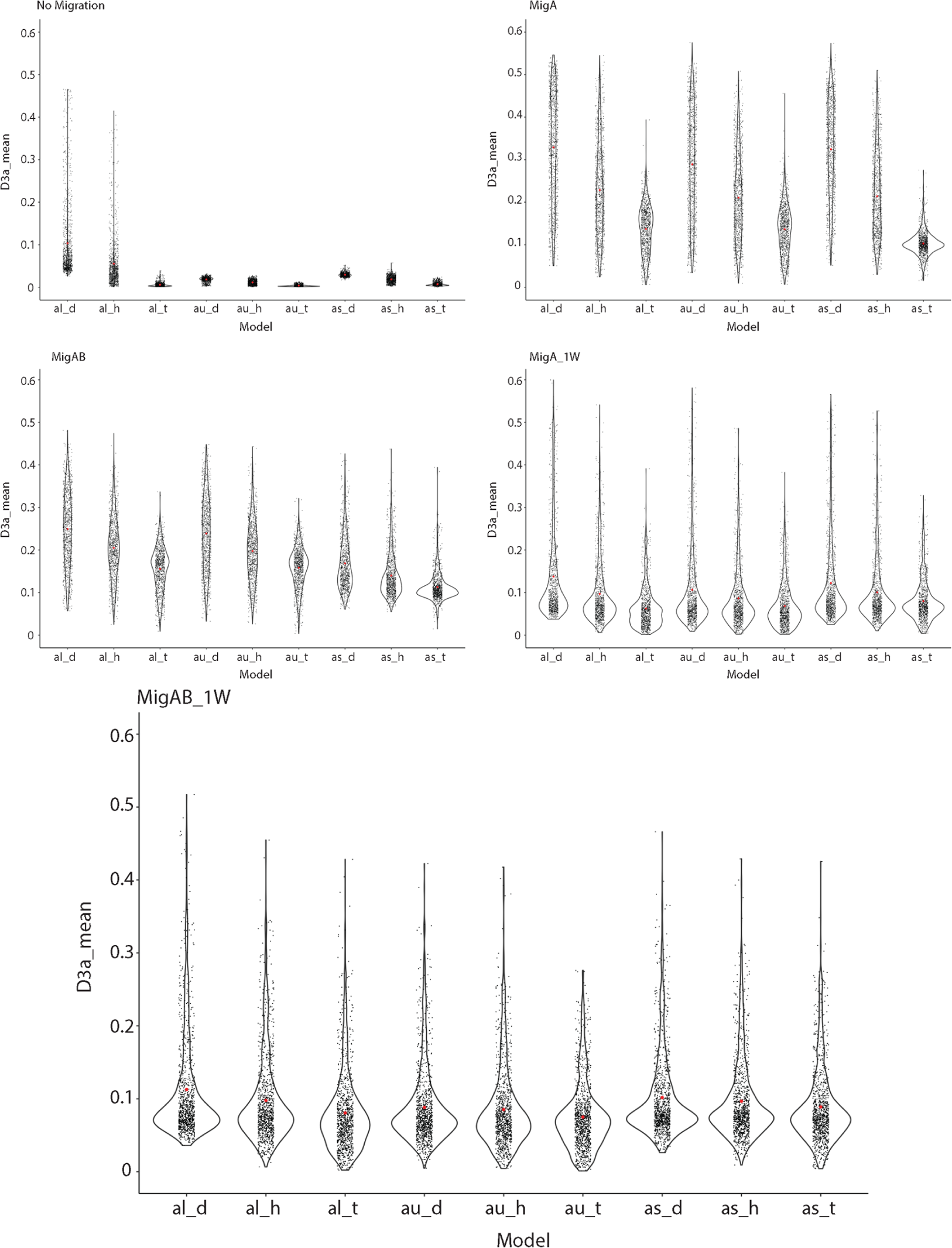
Violin plots and observed values for the average of the D3a statistic under each simulated polyploid speciation, inheritance, and migration models. Red dots are the average values for each model.

**Supplemental Figure 13.**
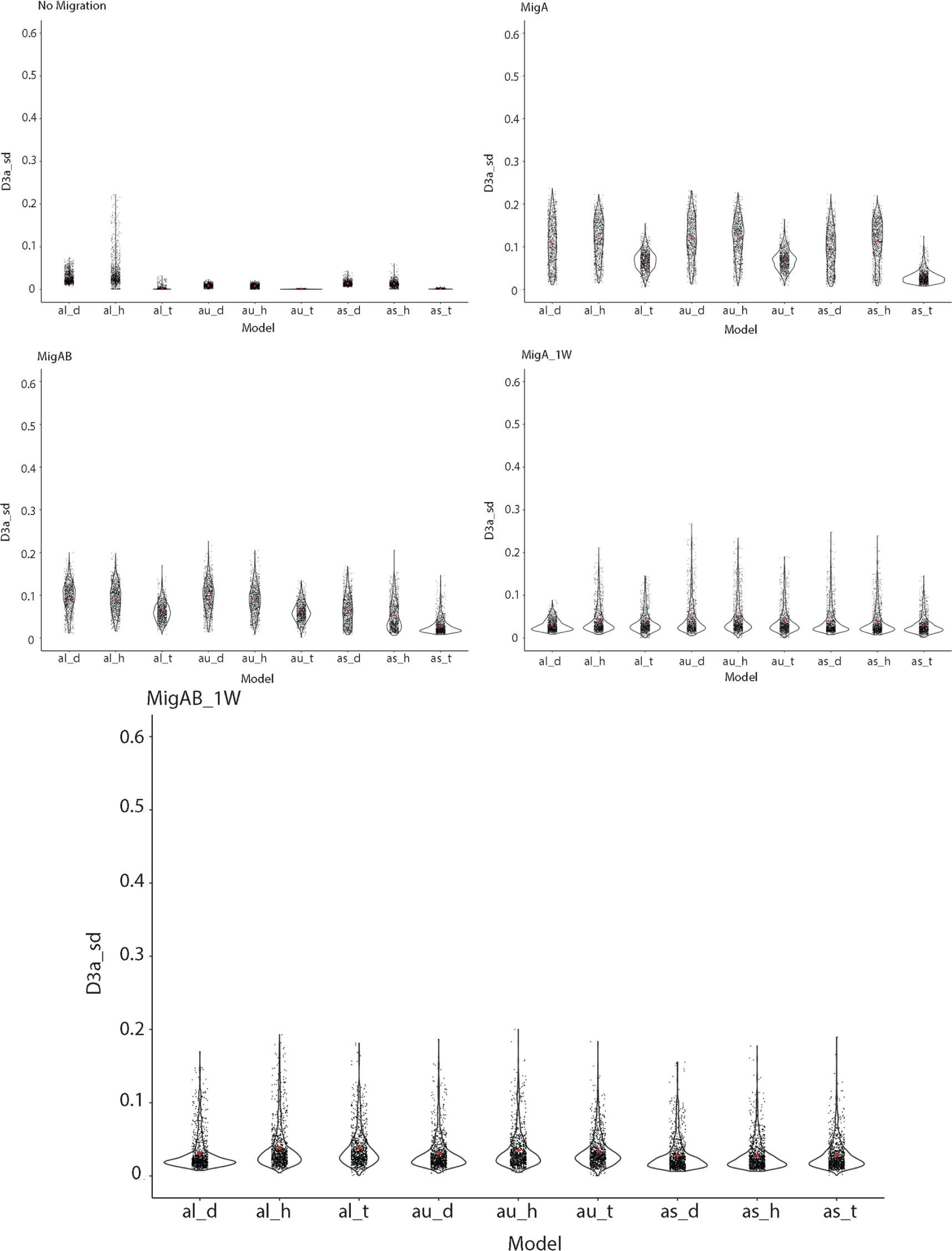
Violin plots and observed values for the standard deviation of the D3a statistic under each simulated polyploid speciation, chromosomal inheritance, and migration models. Red dots are the average values for each model.

## Notes

### Competing Interest Statement

The authors have declared no competing interest.

